# The role of autoproteolysis and mitoribosomal proteins in regulation of mitochondrial LACTB tumor suppressor

**DOI:** 10.64898/2025.12.13.694101

**Authors:** Sara Escudeiro-Lopes, Juan M. Gonzalez-Morena, Alzbeta Baudysova, Michal Svoboda, Pavel Marasek, Gabriel Rodriguez-Gomez, Jessica Ferreira Mendes, Chung Weng Phang, Sravan Kumar Miryala, Pavel Jakoube, Beata Kovarova, Susana Machado, Yuxiong Feng, Zuzana Keckesova

**Affiliations:** Institute of Organic Chemistry and Biochemistry, Czech Academy of Sciences, Prague, Czech Republic; Faculty of Science, Charles University, Albertov 6, 128 43 Prague, Czech Republic; Zhejiang Provincial Key Laboratory of Pancreatic Disease, First Affiliated Hospital, and Institute of Translational Medicine, Zhejiang University School of Medicine, Hangzhou, China; Cancer Center, Zhejiang University, Hangzhou, China

**Keywords:** LACTB, tumor suppressor, autoproteolytic activity, MRPS34, breast cancer

## Abstract

Tumor suppressors represent one of the first lines of defense against malignant transformation and their inactivation in cells leads to onset of tumorigenesis. Cancer cells employ a variety of ways to inactivate cellular tumor suppressors, such as their epigenetic silencing or mutagenesis. Less understood are mechanisms by which cancer cells inactivate tumor suppressors post-translationally. Here, we uncovered a previously undescribed post-translational strategy that cancer cells use to inactivate the potent mitochondrial tumor suppressor LACTB in breast cancers. We discovered that substrate of LACTB can be LACTB itself; that LACTB possesses autoproteolytic ability, which is important for the modulation of its tumor suppressor activity. We show that cancer cells misuse this feature of LACTB to force LACTB into self-degradation. This is mechanistically realized through upregulation of mitochondrial MRPS34 protein, which, through interaction with LACTB, is a positive regulator of the autoproteolytic activity of LACTB and a negative regulator of LACTB. This study, through *in vitro*, *in vivo*, human clinical tumor samples and mutagenesis, provides important new insights into how cancer cells fine-tune the expression and activity of tumor suppressors to promote tumorigenesis.

**Statement of Significance:** We uncovered a unique post-translational strategy and mechanism cancer cells employ to inactivate the potent mitochondrial tumor suppressor LACTB in breast cancers thus expanding our knowledge on regulatory and adaptive mechanisms cancer cells use to silence tumor suppressors.

## Introduction

Deactivation of tumor suppressors is a crucial step in the development and progression of malignancies. In human cells this can occur by complete loss/inactivation of a particular tumor suppressor, through cellular processes such as gene mutation [1,2], aberrant splicing [3,4], and epigenetic silencing [5,6]. The complete loss of tumor suppressor is described by the Knudson’s two-hit hypothesis, which stipulates that inactivation of both alleles of a tumor suppressor gene is required to promote tumor progression [7]. However, subsequent research revealed the existence of a different class of tumor suppressors termed “haploinsufficient/dose-dependent”, where loss of only one allele of such a tumor suppressor already contributes to the tumorigenic process, which is sometimes manifested through lower expression levels of such proteins [8,9]. Examples of happloinsufficient tumor suppressors include p27Kip1, NF1, and PTEN, among others [9]. These types of tumor suppressors represent attractive therapeutic options for novel cancer treatments since, in many cases, they are never completely absent from the cell. Recently, beta-lactamase-like protein (LACTB) was also characterized as a novel happloinsufficient mitochondrial tumor suppressor in breast cancer cells [10].

LACTB is a ubiquitously expressed mitochondrial protein, derived from the bacterial penicillin-binding/β-lactamase proteins (PBP-βL) family, which eukaryotic cells acquired during endosymbiosis [11,12]. Eukaryotic LACTB shares conserved active site motifs with bacterial PBP-βL proteins, including the catalytic serine site -SXXK- [12]. In bacteria, PBP-βL proteins are involved in peptidoglycan synthesis, which is important for the bacterial cell wall biogenesis [13,14]. Since eukaryotic cells lack bacterial peptidoglycans, the need for LACTB’s catalytic activity is currently unknown.

LACTB was first described as a tumor suppressor in breast cancer [10], and follow up studies extended its tumor suppressor role to other cancer types, such as colorectal cancer [15–18], hepatocellular carcinoma [19], glioma [20], gastric cancer [21], melanoma [22,23], lung cancer [24] and ovary cancer [25]. However, studies in pancreatic cancer [26] and in nasopharyngeal carcinoma [27] showed that higher expression of LACTB was associated with cancer progression, thus identifying LACTB as a possible tumor promoter in these cancer types. LACTB’s tumor suppressor function is realized through modulation of lipid metabolism, inhibition of cellular proliferation and epithelial-mesenchymal transition (EMT), and induction of differentiation and cancer cell death, through several different mechanisms of action in a tissue-specific manner [10,15,19,20,22,28]. Collectively, these studies showed that downregulation/inactivation of LACTB in cancer cells which occurs through both, transcriptional and post-translational mechanisms, is a common mechanism how cancer cells counteract the negative effects of LACTB on their viability. On a transcriptional level this regulation happens through different mechanisms: LACTB silencing occurs through binding of microRNAs to the 3’-UTR of LACTB: microRNA-1276 in colon cancer cells [18] and MiR-125b-5p in the context of non-tumorigenic THP-1 macrophages [29]. In colorectal cancer, LACTB downregulation is associated with its promoter methylation and Histone H3 hypoacetylation, showing the involvement of epigenetic modifications in LACTB repression [15]. In melanoma, the transcription factor SOX10 binds to the LACTB promoter and negatively regulates its transcription [22]. However, while some information is known about the transcriptional regulation of LACTB, the mechanisms that lead to post-translational downregulation of LACTB by cancer cells are currently unknown.

It was the intention of our work to uncover this important aspect of post-translational regulation of LACTB in breast cancer cells, which started by our search for the LACTB substrate. Using adapted Proteomic Identification of Cleavage Sites (PICS) assay, we identified that the substrate of LACTB could be LACTB itself; that LACTB possesses autoproteolytic activity. This autoproteolytic activity is important for the tumor suppressive ability of LACTB and is realized through targeting the AAPD and TVGD amino acid sequences within LACTB. The autoproteolytic activity of LACTB is hijacked by cancer cells that use it to efficiently downregulate LACTB tumor suppressor protein. Mechanistically, this is achieved by the upregulation of the 28S ribosomal protein S34 (MRPS34) protein, which acts as a negative regulator of LACTB. Through their interaction, MRPS34 binds LACTB and stimulates its autoproteolytic activity that leads to LACTB’s self-degradation in cancer cells and increased cancer cell proliferation and tumor growth.

## Material and Methods

### Cell culture and reagents

Cells were cultivated at 37 °C in 5 % CO_2_ humidified atmosphere. MCF-7 were purchased from Merck and modified by introduction of H-RAS V12 oncogene to form estrogen-independent MCF7ras (MCF7R) cells. HMEC cells were purchased from Lonza and were immortalized by the introduction of H-TERT to form HME cells. HME cells were grown in a 1:1 mixture of DMEM/Nutrient Mixture F-12 Ham (Sigma-Aldrich) and Mammary Epithelial Cell Growth Basal Medium (MEBM, Lonza) supplemented with Mammary Epithelial Cell Growth Medium BulletKit™ (MEGM™, Lonza). Cell lines, growth medium and sources are described in Supplementary Table 1. All cell lines used in this study were verified to be mycoplasma negative. The MG132 was purchased from Sigma-Aldrich (M8699) and used at 20 µM final concentration.

### Constructs

FUW-LACTB-Flag was derived from FUW-LACTB vector (a kind gift of Dr. Robert Weinberg, MIT) with the introduction of Flag-His tags to the C-terminus of LACTB. pCLNCX-LACTB-Flag-His was a kind gift from Daniel Bachovchin (Sloan Kettering Institute, New York, USA). FUW-LACTB-AAPA+TVGA-Flag-His was generated by mutating the aspartic acids into alanine in the LACTB aminoacid residues 73 and 406. FUW-LACTB-S164A and pCLNCX-LACTB-S164A-Flag-His were prepared by mutating the serine at position 164 to alanine. Mutagenesis was performed using Quick Change XL Site-Directed Mutagenesis Kit (Agilent Technologies), according to manufacturer’s protocol. pLV-shMRPS34-EGFP-blast (vector ID: VB900084-4794rbw) and doxycycline-inducible pLV-TRE-MRPS34-blast (NM_001300900.2) were purchased from VectorBuilder. Human MRPS34 cDNA was purchased from Vector Builder (hMRPS34 [NM_001300900.2]) and sub-cloned into PLVX-puro vector through restriction enzymes BamHI and EcoR1, and a Flag-His-tag was inserted at MRPS34 C-terminus. The sequence of the primers used for mutagenesis, cloning and sequencing are described in Supplementary Table 2.

The doxycycline inducible system used throughout this study consists of two separate vectors: one vector carrying the reverse tetracycline-controlled transactivator (rtTA) and second vector carrying the Tet responsive element followed by cloned gene of interest. For activation of the tetracycline-inducible gene expression, the cells were treated with 1 μg/ml of doxycycline hyclate (DOX, Sigma-Aldrich) in the medium. To avoid non-specific DOX effects, our control cells were also treated with DOX, where applicable.

### Lentiviral vector preparation and transduction

40% confluent HEK293T cells were transfected with 1 μg of pMDG, 1 μg of pCMV delta R8.2 (Addgene plasmid # 12263) and 1.5 μg of the gene-encoding vector (such as PLVX-puro, FUW-puro or pLV-blast) in DMEM 10% FBS without antibiotics using X-treme gene HP DNA transfection reagent (Roche) mixed in Opti-MEM (Gibco). Medium on transfected cells was changed after 24 hours, and lentiviral vector particles were collected at 48- and 72- hours post transfection. Viral particles were filtered using 0.45 μm filter.

For lentiviral transduction, cells were seeded at 30 % confluency in a 10-cm dish and transduced 24 h later with virus in the presence of 5 μg/ml Polybrene (Merck). Cells were then selected by the relevant selection marker.

### Cell proliferation assay

For cell proliferation assessment, 3.0x10^5^ cells were seeded in a 10-cm plate, unless stated otherwise. 24-hours later, after cell attachment, they were treated with 1 μg/ml of doxycycline hyclate (DOX, Sigma-Aldrich). Live cells were measured after 3, 6, 9 and 12 days of DOX treatment in an automatic cell counter (Nano EnTek) using a 1:1 ratio of Trypan blue and medium containing the trypsinized and well homogenized cells. Fresh DOX was added every 3 days.

### Recombinant protein preparation

HEK293T cells expressing pCLNCX-LACTB-Flag-His were grown to 90 % confluency on twenty 15-cm plates. Cells were washed twice with PBS and scraped. Cell pellets were then collected by centrifugation at 3000 x g for 5 minutes. The pellets were re-suspended in cell lysis buffer containing 20 mM TRIS; 150 mM NaCl; 1 mM EDTA; 1mM EGTA; 1 % igepal; 1 % sodium deoxycholate; pH 7.5; debris was removed by centrifugation at 12000g for 15 minutes. LACTB was then captured by incubation of the cell lysate with anti-Flag M2 Magnetic Beads (Sigma). Protein lysate and beads rotated for 2 hours or overnight at 4 °C. Afterwards, beads were washed three times with PBS and 150 µg of 3x Flag Peptide (Sigma) was added to the beads for 1 hour at 4 °C, for protein elution. Beads were removed and the supernatant containing the recombinant protein was concentrated using an Amicon Ultra Centrifugal Filter with 30 kDa cut-off (Milipore). Final concentration of recombinant protein was determined by NanoDrop microvolume spectrophotometer. LACTB purified proteins were kept non-frozen at 4 °C until used for the experiment. Before the subsequent experiments, the quality of LACTB recombinant protein was always validated by enzymatic cleavage assay to ensure the functional catalytic activity of LACTB. On rare occasions when the catalytic activity of isolated LACTB was inadequate, the recombinant protein was discarded. Recombinant proteins for MRPS34 (TP300834), GAPDH (TP302309), COQ6 (TP760736), HSBP1 (TP301399), ASCL3 (TP320042) and TOMM20 (TP310746) were purchased from OriGene. Recombinant protein for HIGD1A (ab125988) was purchased from Abcam. Proteins were aliquoted and stored at -80°C.

### LACTB *in vitro* substrate assay

150 ng of recombinant protein was added to a Nunc MicroWell 96-well black, clear optical-bottom plates with polymer base (Sigma-Aldrich) in PBS, together with 1 µl of peptide substrate Ac/nonAc-xxxD-AMC (Enzo Life Sciences and IOCB peptide synthesis facility) (prepared in DMSO at 10 mM) at a final volume of 100 μl. Where needed experiments were performed with Granzyme B inhibitor (Sigma-Aldrich, 368050) or PBS was substituted with H2O. Fluorescence was measured on Spark Multimode Plate Reader (Tecan) for 50- to 180- minutes at 25°C with a kinetic cycle of 10 minutes, fluorescent top reading of multiple reads per well and with an excitation wavelength of 380 nm and an emission wavelength of 460 nm. The peptide substrates used contained aspartic acids present within the LACTB sequence (Supplementary Table 3); as a negative control we used a tetrapeptide ending in alanine instead of aspartic acid: Ac-YGAA-AMC; as a positive control we used a tetrapeptide that has already been described to be cleaved by LACTB: Ac-YVAD-AMC [10].

### LACTB autoproteolytic activity assay

Purified recombinant LACTB in H_2_O (distilled and autoclaved) or PBS was incubated at 25 °C, with and without Z-AAD-CMK (Granzyme B inhibitor), which was already described to inhibit LACTB activity [30], for different timepoints. Samples were then processed for Western Blot analysis and immunoblotted for LACTB (as described below). Of note, the exact timing of the occurrence of the autoproteolytic activity depends on the quality and purity of the recombinant LACTB and on the duration of its storage before the onset of the experiment. Therefore, based on these parameters it sometimes manifests several hours earlier or later than the average times shown in this manuscript.

### Modified PICS assay

Mitochondria were isolated from twelve 15-cm plates of confluent MCF7R cells. Mitochondria isolation was done using Qproteome Mitochondria Isolation Kit (Qiagen), according to manufacturer’s instructions. Isolated mitochondria were dissolved in 1 ml of Outer Mitochondrial Membrane Disruption Buffer (10 mM HEPES and 1 mM EDTA, pH 7.4) and incubated for 30 minutes at 4 °C. Then 50 µl of 3 M KCl (stock concentration) was added and incubated for 10 minutes at 4 °C. After 10 minutes sample was spin down at 11000 x g for 10 minutes, the supernatant was sonicated and the pellet was dissolved in 1 ml of Inner Mitochondrial Membrane Disruption Buffer (5 mM HEPES, 5 mM sucrose and 1 mM EGTA, pH 7.2), incubated for 30 minutes at 4 °C and then sonicated. To both sonicated samples RQ1 DNase (Promega) was added for 30 minutes at 25 °C. The mixture was then centrifuged at 11000 x g, pellets were kept for later use and the supernatant was concentrated to 200 µl final volume using Amicon filter with 3 kDa cut-off (Millipore). Protein concentration in the sample was then determined by DC Protein Assay (BioRad). Sulphate groups were blocked using 5 mM of dithiothreitol (DTT) that was added to the isolated mitochondrial proteome sample and incubated at 37 °C for 1 hour. This was followed by the addition of 40 mM iodoacetamide and incubation at 37 °C for 1.5 hours. The reaction was terminated with addition of another 15 mM DTT and incubation for 10 minutes at 37 °C. Afterwards, amino groups were blocked with 60 mM formaldehyde and 60 mM sodium cyanoborohydride for 18 hours at 37 °C. The reaction was stopped by addition of 35 mM glycine. The final sample was washed 15 times with PBS using amicon filter with 3 kDa cut-off (Millipore) to reduce the concentration of glycine. Recombinant LACTB-WT or LACTB-S164A was added to the sample of mitochondrial proteins with blocked reactive groups. The addition was done in two steps – first, 0.5 µg of LACTB was added and mixture was left for 2 hours at 25 °C. Then another 0.5 µg of LACTB was added and mixture was incubated for 2 hours at 37°C. To trap newly formed primary amines, sample was incubated with 0.5 µl of 10 mM Dde Biotin-PEG4-NHS ester (Click Chemistry Tools) solution in DMSO for 2 hours at 25°C. Biotinylated proteins were then purified by the addition of Streptavidin Magnetic Beads (Dynabeads™ M-280 Streptavidin, Invitrogen) and kept at 25°C for 30 minutes with mild agitation. Beads were then collected, washed thoroughly with 1 ml of different solutions in the following order – PBS, 1 M guanidin HCl, PBS, 20 % isopropanol, PBS, 5 % DMSO, PBS, 5 % ACN and ten times beads equilibration buffer (50 mM HEPES and 150 mM NaCl, pH 7.5). Bound proteins were then eluted using 2 % hydrazine in 100 mM sodium phosphate and analysed by mass spectrometry.

### Immunoprecipitation (IP) assay

IP was done with crosslinking and non-crosslinking conditions. MCF7R-RTTA-hygro-FUW-LACTB-WT (MCF7R-LACTB-WT) cells were treated with DOX for 1 day and then washed twice with PBS. For crosslinking conditions: 1mM of DSP (Thermo Scientific) in PBS was added to the cells for 20 minutes at room temperature, followed by 10 mM Tris pH 7.5 for 15 minutes at room temperature in order to stop the DSP reaction. Cells were lysed in IP-lysis buffer containing 20 mM Tris, 150 mM NaCl, 1mM EDTA, 1 mM EGTA, 0.5 % Triton x-100, and supplemented with protease (cOmplete, Roche Diagnostics) and phosphatase (PhoStop, Roche Diagnostics) inhibitors. Cell lysates were sonicated three times for 10 seconds and centrifuged at 12000g for 15 minutes to remove cell debris. Protein lysates were quantified using DC protein assay kit (BioRad). 2 μg of antibody was added to 50 μl of protein G magnetic Dynabeads (50% slurry) previously washed in IP-lysis buffer. Beads with antibody were rotating for 2 hours at 4 °C. For crosslinking conditions: 20 mM of DMP (Thermo Scientific) in sodium borate pH 9 was added to the beads, rotating for 30 minutes at 25 °C, process repeated twice; 50 mM glycine pH 2.5 was used to stop the DMP reaction. 1.5 mg of protein lysate were added to the beads pre-washed with IP-lysis buffer. Lysates with beads were rotating for at least 2 hours at 4 °C. Protein was eluted from the beads by boiling for 10 minutes at 98 °C with Laemmli buffer (BioRad) and reducing agent (Invitrogen). Co-IP for LC-MS/MS analysis: beads were rinsed 3x with 20 mM Tris pH 8.0 to remove salts and detergent and kept in this buffer until the analysis. Bound proteins (LACTB and co-immunoprecipitated proteins) were analyzed via LC-MS/MS after in-solution trypsin digestion. Beads with no loaded antibody were used as a control for unspecific interactions. Each condition was prepared in triplicates. Proteins detected in LACTB co-IP replicates and undetected in the control sample were considered as positive hits.

### Quantitative RT-PCR

Total RNA was isolated directly from cultured cells using the RNeasy Plus Mini kit (Qiagen), following the manufacturer’s instructions. Reverse transcription was performed with a High Capacity cDNA Reverse Transcription Kit (Life Technologies). mRNA levels were measured with gene-specific primers using the SYBR Green I master mix (Roche) in a Roche LightCycler 480 system (Roche), using a 384-well plate; each well consisted of 25 ng/ml of cDNA. The amplification program consisted of an initial cycle of 95°C for 15 s, denaturation at 95°C, followed by 45 cycles of 60°C. After amplification runs, quantification cycle (Cq) values were provided by Roche LightCycler 480 system. Cq value of each sample was normalized using the geometric mean Cq value of the reference gene, corresponding to DCT. Then, the DDCT was calculated, which refers to the Cq of the sample of interest. Lastly, the value of 2-DDCt was considered. The obtained data were then further normalized to the expression of these genes relative to the control cells. GAPDH was used as reference gene for normalization. The PCR primer sequences are listed in Supplementary Table 2.

### Western blot analysis

Proteins were extracted from cells using RIPA buffer (Sigma) supplemented with protease and phosphatase inhibitors. Approximately 20 μg of protein lysate was loaded into SDS-PAGE 4 % to 15 % gradient gels (Bio-Rad). Western blots and transfers were done following standard protocols in a wet-transfer system (Bio-Rad) to PVDF membranes (Immobilon-P, Millipore). Next, membranes were blocked for 1 hour with 5 % non-fat milk in PBS with 0,1 % Tween-20 (v/v) followed by blotting with relevant antibodies overnight at 4°C. Horseradish peroxidase-conjugated secondary antibodies were used (Cell Signaling, 1:5000). Blots were developed using enhanced chemiluminescence (Dura or Femto, Thermo Scientific), signal was detected using Azure c600 Western blot Imaging system. Antibodies used are described in Supplementary Table 4.

### Dot blot analysis

PVDF membranes were activated in methanol and placed on the top of a marked grid Whatman filter paper soaked in transblotting buffer. Recombinant proteins were spotted into the wet membrane and incubated for 15 minutes to fix the proteins. Membrane was then blocked for 1 hour in 5 % non-fat milk in PBS with 0,1 % Tween-20 (v/v), followed by 2 hours room temperature incubation with recombinant LACTB (around 10 μg) in PBS. The membrane was washed 3 times in PBS-T and blotted overnight at 4 °C using LACTB antibody.

### *In vivo* mouse experiments

All manipulations with cells and mice were performed in Class II Biological Safety Cabinets to achieve aseptic working environment during the whole study. Cells were cultivated in media (DMEM) + 10 % FBS + 1 % Penicilin-streptomycin and suspended in 50% Matrigel prior to injections. Adult female mice strain NSG (NOD.Cg-Prkdcscid Il2rgtm1Wjl/SzJ) were obtained from The Jackson Laboratory. The injected number of MCF7R cells was 3x10^5^ and of HCC1806 was 1,5x10^5^, prepared in 15-30 μL for each dose applied into mammary fat pad bilaterally. Body condition score and tumor size were measured twice weekly. The *in vivo* DOX treatment was administered through drinking water containing 2 mg/ml of DOX and 10 mg/ml of sucrose when tumors reached approximately 4 mm in diameter. The mice were euthanized when the tumors reached cumulative size of 1,5 cm or earlier. Sample sizes were chosen to reach statistical significance, and tumor measurements and data analysis were performed in a blinded fashion. The animal experiment was approved by the Animal Research Ethics Committee of Czech Academy of Science approval ID: AVCR 5282/2021 SOV II.

### Immunofluorescence (IF) assay

LACTB-WT and LACTB-AAPA+TVGA overexpression was induced in MCF7R cells for 1 day, in 13mm glass coverslips. 24 hours after, cells were washed 2 times in PBS for 4 minutes and subsequently fixed with 4 % paraformaldehyde (PFA) and permeabilized with 0.1 % Triton X-100 in PBS, for 20 minutes. Cells were washed 3 more times with PBS for 10 minutes and blocked with 5 % BSA in PBS for 30 minutes. After this step, the coverslips were transferred to a wet chamber, where 50 μl of the diluted primary antibody were added on the top of each coverslip, for at least 1 hour. Afterward, the primary antibody was removed, and cells were washed 3 times with PBS-0.1 % Tween (PBS-T), followed by the addition of the Alexa Fluor secondary fluorescent antibodies for 1 hour. Cells were then washed 3 times with PBS-T and coverslips were mounted in ProLong Gold anti-fade reagent with DAPI (Life Technologies). All steps were performed at room temperature. Antibodies used are described in Supplementary Table 4.

Images were acquired in Zeiss LSM 980 confocal microscope with 405 nm and 561 nm laser excitation wavelengths using 63× (NA 1.4) immersion oil objective. Images correspond to one channel and one section in the z axis in the middle of the cell. Images were analyzed with Image J. Antibodies used are described in Supplementary Table 4.

### Proximity Ligation Assay (PLA)

To evaluate the interaction between endogenous LACTB and the potential binding partner proteins *in situ,* we used the Duolink Proximaty Ligation Assay (PLA) (Sigma-Aldrich) in MCF7R control and MCF7R-FUW-LACTB-WT following the manufacturer’s instructions. Briefly, cells were grown on eight well chamber slides (Lab-Tek Chamber Slide, 177445). For mitochondrial staining, cells were incubated with MitoTracker Deep Red (Thermo Fisher, M22426) at a final concentration of 0.25 µM for 40 minutes at 37°C. The cells were fixed using prewarmed 4,2 % PFA, permeabilized with 0,1 % Triton X-100, incubated with blocking solution (provided by the kit) for 1 hour at 37°C, and then incubated in a solution containing the primary antibodies (LACTB and the potential binding partner/control protein) overnight. PLA probes were prepared by 1:5 dilution in antibody diluent (provided by the kit), added to the slides, and incubated for 1 hour at 37 °C. The ligation of the PLA probes was performed by incubating the cells in the ligation solution for 30 min at 37 °C followed by an amplification step for 100 min at 30 °C. Then the slides were let dry in the dark at room temperature and mounted with Duolink *In Situ* Mounting Medium with DAPI (Sigma-Aldrich). Images were acquired in a confocal microscope (Zeiss) as described above. Quantification of PLA dots was done following the method described in [31].

### Immunohistochemistry (IHC)

Human tumor and normal tissue sections (obtained from TissueArray; BR1191) were deparaffinized by heating the slides at 60 °C for 10 minutes. Next, the slides were washed twice in xylene for 5 minutes, and 1 time in 1:1 xylene:ethanol for 3 minutes. Then slides were hydrated in washes with 100 % (twice), 95 %, 80 %, 75 % and 50 % ethanol for 3 minutes each of them, followed by wash with 0.85 % NaCl, PBS and H_2_O for 5 minutes each. Tissues were boiled for 20 minutes in 1x citrate buffer (Sigma-Aldrich). Slides were washed in PBS for 1 min, and then permeabilized with 0,2 % Triton X-100 for 20 minutes. The blocking solution was added for 20 min (30 % horse serum in PBS, 0,1 % Tween-20 at room temperature). Sections were incubated at 4 °C overnight with primary antibody (antibodies are described in Supplementary Table 4). After three washes with PBS, sections were incubated with secondary antibodies (VECTASTAIN ABC-AP) for 1 hour at room temperature, washed three times with PBS. Then the slides were incubated with solution Fast Red (Abcam). Hematoxylin was used as counterstain to stain nucleus. Slides were dehydrated and mounted. IHC images were acquired in optical light microscope (Nikon).

### EdU staining

MCF7R cells overexpressing LACTB, MRPS34 and LACTB+MRPS34 for different time points were collected. Cells were treated with 20 μM EdU for 90 minutes at the end of LACTB/MRPS34 induction. 600.000 cells were collected for all the conditions. Edu labeling was performed for 30 minutes using 2 mM CuSO_4_, 8 μM CY-5 (Sigma) and 20 mg/ml ascorbic acid. Cells were also stained with 3 μg/ml of DAPI for 15 minutes to stain the DNA. The signal was detected by flow cytometry and data analyzed using FlowJo 10.7.1 software.

### Statistical analysis

Data is shown as the average ± standard deviation (SD). Comparative studies between quantitative variables following normal distribution were evaluated using Student’s *t*-test for paired or unpaired samples according to the experiment. For *in vivo* experiments differences in tumor growth were analyzed by 2-way ANOVA. Differences were considered significant when p value ≤ 0.05. Statistical analysis was performed using GraphPad Prism 8 software.

## Results

### 1. LACTB possesses autoproteolytic activity

To identify the substrate of LACTB we set up and optimized a modified protocol of the Proteomic Identification of Cleavage Sites (PICS) assay [32] for uncovering substrates of proteases (Fig. 1A). In this approach, we utilized the MCF7ras cell line (MCF7R) whose growth, as shown by us previously, is negatively affected by LACTB-WT induction, but not by the catalytic mutant LACTB, implying the presence of LACTB substrate in this cell line [10]. We isolated full-length proteome of mitochondria-enriched fraction from MCF7R cells, which was followed by blocking the reactive groups within the proteome and addition of wild type recombinant LACTB. Cleavage of its substrate by LACTB revealed new reactive N-termini within the cleaved substrate, which were biotinylated, captured and analyzed by mass spectrometry. In parallel, the assay was performed under the same conditions in the presence of catalytically inactive S164A LACTB mutant as a negative control. The modified assay was performed in two biological replicates (2 biological replicates for LACTB-WT and 2 biological replicates for S164A LACTB). The results identified 15 proteins with a high probability of being the substrate/s of LACTB (Fig. 1B). The recombinant LACTB protein used in the modified PICS assay was isolated from human HEK293T cells and validated by Western blot analysis and by enzymatic activity assay for the correct size, expression, and functional catalytic activity (Suppl. Fig. 1A-C).

**Figure 1.**
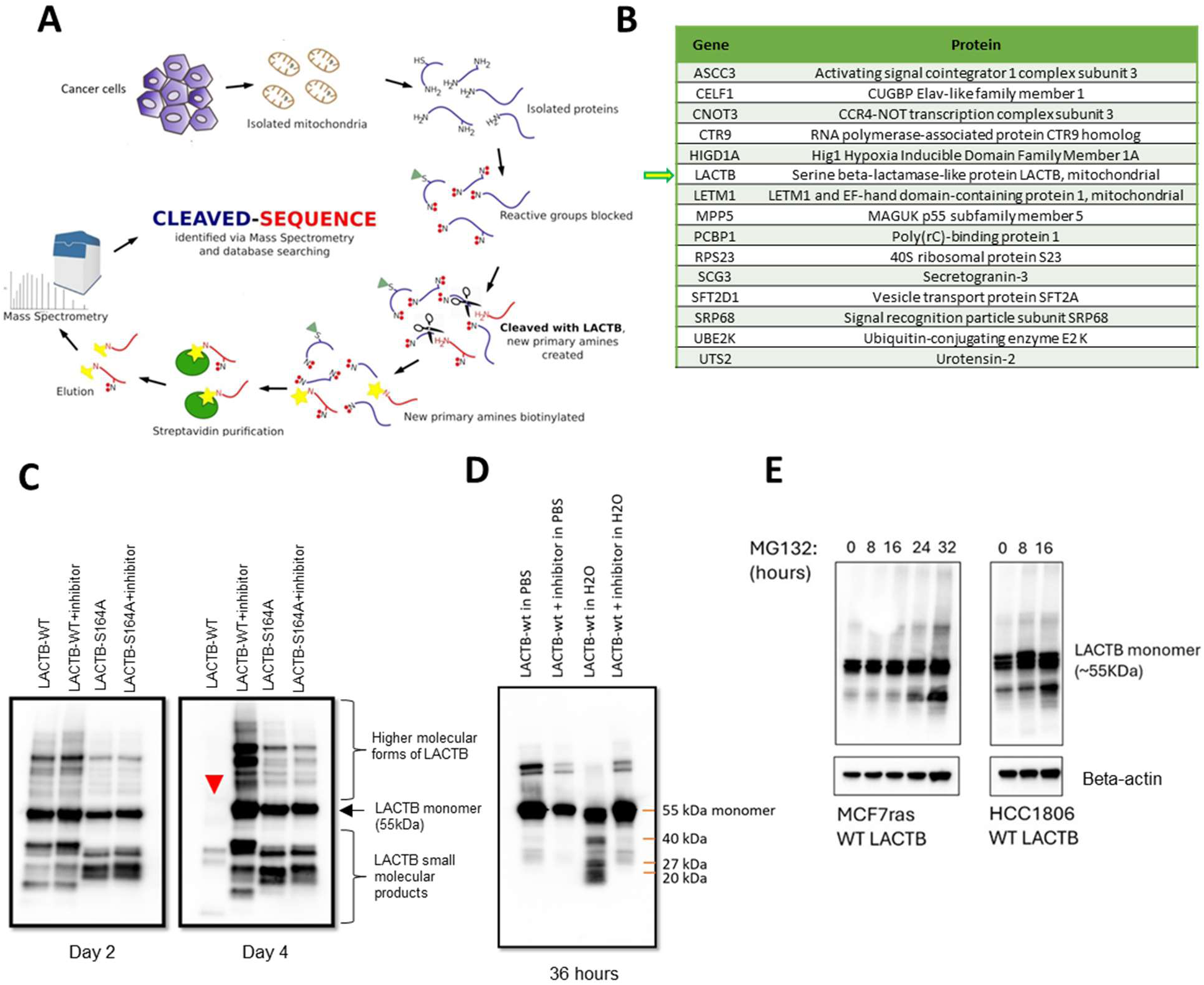
Identification of LACTB substrates. **(A)** Schematic representation of the workflow of the modified PICS assay. Assay was performed with both, wt LACTB and S164A LACTB. **(B)** Table summarizing the potential LACTB substrates. **(C)** Recombinant LACTB-WT and S164A LACTB, with and without an inhibitor of its activity (Z-AAD-CMK) were incubated at different time points at 25 °C and visualized by Western blot. The experiment was performed twice. Red arrow points to the disappearance of the LACTB monomer. **(D)** LACTB-WT, with and without inhibitor, incubated either in PBS or in H_2_O for 36 hours at 25 °C. The experiment was performed three times. **(E)** Western blot analysis of LACTB-WT (induced by doxycycline for 24 hours) in MCF7R and HCC1806 cells upon treatment with 20 µM MG132. Membrane was probed with LACTB and beta-actin antibodies.

Once the list of potential LACTB substrates was generated, we were surprised to notice that LACTB-WT was amongst the substrate hits (Fig. 1B and Suppl. Data 1). Therefore, in this manuscript, we set out to investigate the possibility of LACTB serving as its own substrate and possessing autoproteolytic activity. We incubated recombinant LACTB-WT and LACTB-S164A, with and without the inhibitor of its enzymatic activity (Granzyme B inhibitor - Z-AAD-CMK, [30]), for 2 days and 4 days in PBS buffer and examined the resulting stability of LACTB by Western Blot (Fig. 1C). The results showed that LACTB-WT was efficiently degraded at 4 days of incubation. This degradation is completely abolished in the presence of the inhibitor of LACTB’s enzymatic activity, where LACTB protein levels did not change at 4 days of incubation. The same result as with the chemical inhibitor of LACTB’s catalytic activity was also observed with genetic S164A LACTB catalytically inactive mutant, whose protein levels were also stable at 4 days incubation time. These data showed that LACTB-WT indeed possesses autoproteolytic activity, which is realized through the active catalytic serine residue in the amino acid position 164. No differences in LACTB-WT stability were seen in earlier time point (2 days) suggesting that while LACTB is capable of autoproteolytic activity, this activity occurs with slow kinetics or might require special cellular conditions, cofactors, or additional stimuli for efficient execution. Interestingly, LACTB’s autoproteolytic activity manifested at earlier time points (36 hours) in H_2_O, as opposed to PBS (Fig. 1D). Next, we examined whether the autoproteolytic ability of LACTB-WT manifests itself also in a physiological cellular context. Upon treatment of MCF7R and HCC1806 cells with MG132 proteasome inhibitor we observed increased accumulation of the LACTB-WT degradation products (Fig. 1E) confirming the ongoing autoproteolytic activity of LACTB in these cancer cell lines.

### 2. The autoproteolytic activity of LACTB is targeting the AAPD/TVGD amino acid residues

We wanted to examine the autoproteolytic activity of LACTB in more detail and to map the amino acid residues within LACTB targeted in this process. Since LACTB was shown to cleave peptides after aspartic acid residues [10,30], we synthesized all D-end tetrapeptides from LACTB sequence (Fig. 2A, Suppl. Fig. 2A) fused with fluorescent AMC-tag and incubated these peptides in the presence of LACTB-WT protein. AAPD and TVGD peptides were most efficiently cleaved by LACTB-WT, suggesting that these two sites might be preferentially targeted in the LACTB autoproteolytic process (Fig. 2B). AAPD is located in a structurally disordered N-terminal LACTB region, and both, TVGD and the catalytic pocket (constituted by SISK, YST and HTG motives), are located on the surface of the LACTB molecule, which would make AAPD and TVGD sites accessible for cleavage (Fig. 2C). Regarding their charge, both AAPD and TVGD are negatively charged, as opposed to the catalytic core which is positively charged (Suppl. Fig 2B) further supporting the notion of their possible interaction. To confirm the role of AAPD and TVGD sites as peptide sites targeted in the autoproteolytic activity of LACTB, we prepared a mutant of LACTB where aspartic acids in these two sites were mutated into alanines (LACTB-AAPA+TVGA). Our experiments showed that, as opposed to LACTB-WT, when the recombinant LACTB-AAPA+TVGA protein was incubated *in vitro*, it lacked the ability of autoproteolytic cleavage (as shown by the complete lack of any degradation products), thus confirming the important role of these sites in the autoproteolytic process (Fig. 2D). Although the autoproteolytic activity is impaired in LACTB-AAPA+TVGA, this non-cleavable mutant still retains its enzymatic activity directed against a peptide probe (Suppl. Fig. 2C), suggesting that the enzymatic activity of LACTB can be directed against external peptide or against LACTB itself independently. We next researched the behavior of this mutant in the cellular assays and expressed doxycycline inducible LACTB-WT and LACTB-AAPA+TVGA for one day in MCF7R and HCC1806 cells and examined by Western blot the appearance of the degradation products of LACTB. Indeed, expression of LACTB-WT led to appearance of robust degradation products while this was not observed in case of the expression of the non-cleavable LACTB-AAPA+TVGA mutant (Fig. 2E). The faint degradation bands present in the LACTB-AAPA+TVGA mutant samples are the result of the endogenous LACTB-WT that is present in the small amounts in both MCF7R and HCC1806 cells.

**Figure 2.**
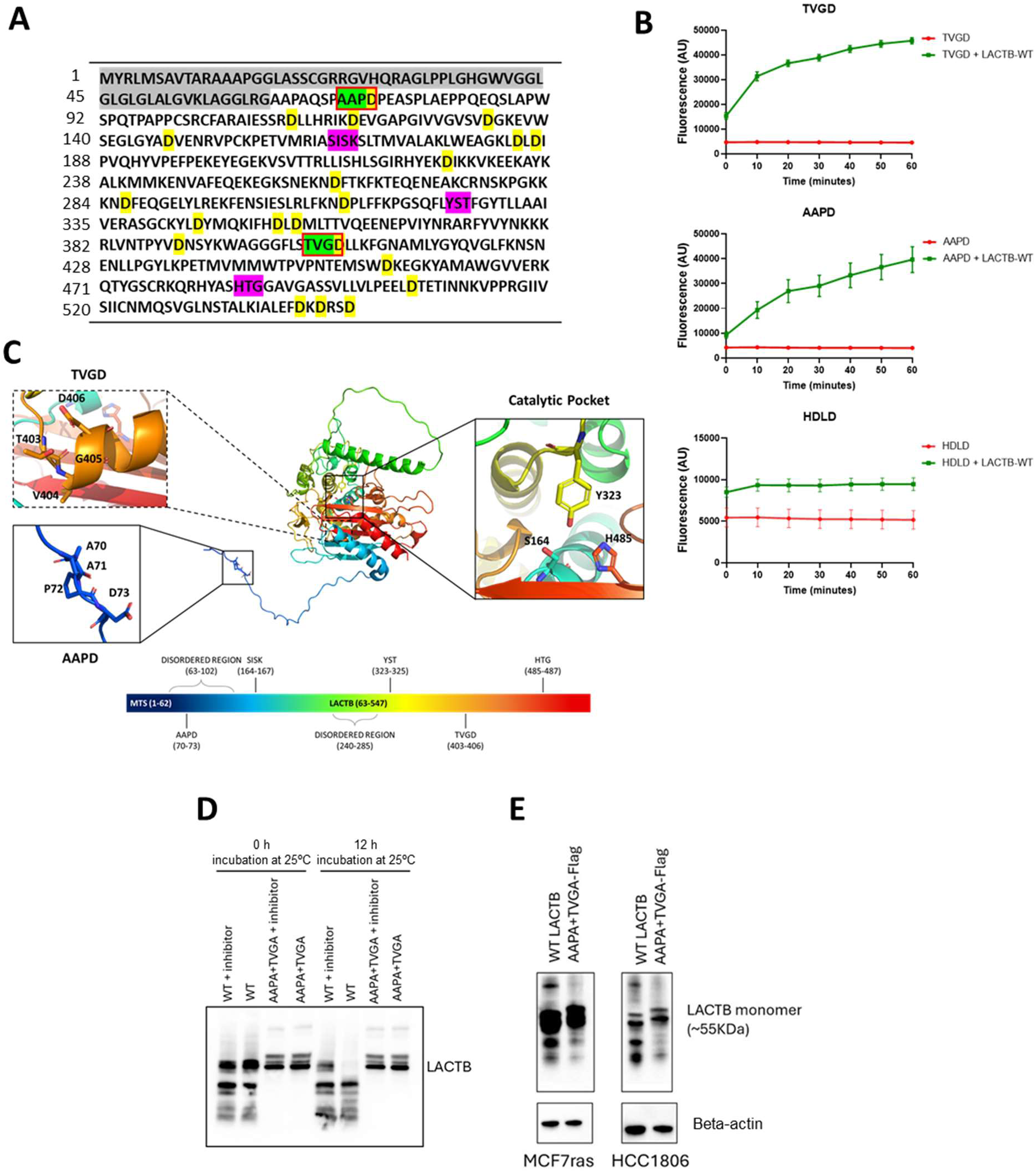
The effect of AAPD/TVGD residues on the autoproteolytic activity of LACTB. **(A)** Amino acid sequence of LACTB-WT. Grey – Mitochondria targeting sequence; Green and red – Sites cleaved by LACTB; Pink – Residues that constitute LACTB’s catalytic pocket; Yellow – aspartic acids within LACTB. **(B)** Enzymatic activity of LACTB-WT against the two top peptide hits – AAPD and TVGD and against a peptide (HDLD) that did not score as a hit. The experiment was performed three times. **(C)** Schematic representation of full length LACTB monomer structure, highlighting the locations of the catalytic pocket, as well as the targeted cleavage sites AAPD and TVGD. **(D)** Western blot analysis of recombinant LACTB-WT and recombinant LACTB-AAPA+TVGA incubated at 25 °C, in H_2_O, with or without the inhibitor Z-AAD-CMK. **(E)** Western blot analysis of LACTB-WT and LACTB-AAPA+TVGA-Flag (both induced by doxycycline for 24 hours) in MCF7R and HCC1806 cells. Membrane was probed with LACTB and beta-actin antibodies.

### 3. The autoproteolytic activity of LACTB influences its tumor suppressive ability *in vitro* and in vivo

We examined whether the autoproteolytic activity of LACTB has any physiological significance and can modulate the tumor suppressive ability of LACTB. We therefore expressed the non-cleavable LACTB-AAPA+TVGA and LACTB-WT in MCF7R and HCC1806 cells. Using Western blot analysis and confocal microscopy, we confirmed that LACTB-AAPA+TVGA mutant is stably expressed in cells, is correctly localized in mitochondria and its levels of expression are comparable to that of LACTB-WT in HCC1806 cells and little decreased in MCF7R cells (Suppl. Fig. 3A-B). We then monitored the growth of MCF7R and HCC1806 cells upon induction of these proteins for up to 12 days. Interestingly, the non-cleavable LACTB-AAPA+TVGA mutant was even more potent in negatively affecting the growth of cancer cells than its wild-type counterpart in both cell types (Fig. 3A). We then proceeded to investigate whether the expression of LACTB-AAPA+TVGA has a negative effect on tumor growth. Control MCF7R cells and MCF7R cells transduced with dox-inducible LACTB-WT or with dox-inducible LACTB-AAPA+TVGA were injected orthotopically into mice and the growth of the tumors was monitored after the induction of both proteins. Tumors in which non-cleavable LACTB-AAPA+TVGA had been induced decreased in size even more significantly than tumors with LACTB-WT (Fig. 3B-C), confirming the role of autoproteolytic activity in LACTB’s tumor suppressive effects. Similar results were observed with HCC1806 tumors (Suppl. Fig. 3C-D). These results suggest that LACTB protein impaired in its autoproteolytic ability is even more potent in negatively affecting cancer cells *in vitro* and *in vivo* than LACTB-WT.

**Figure 3.**
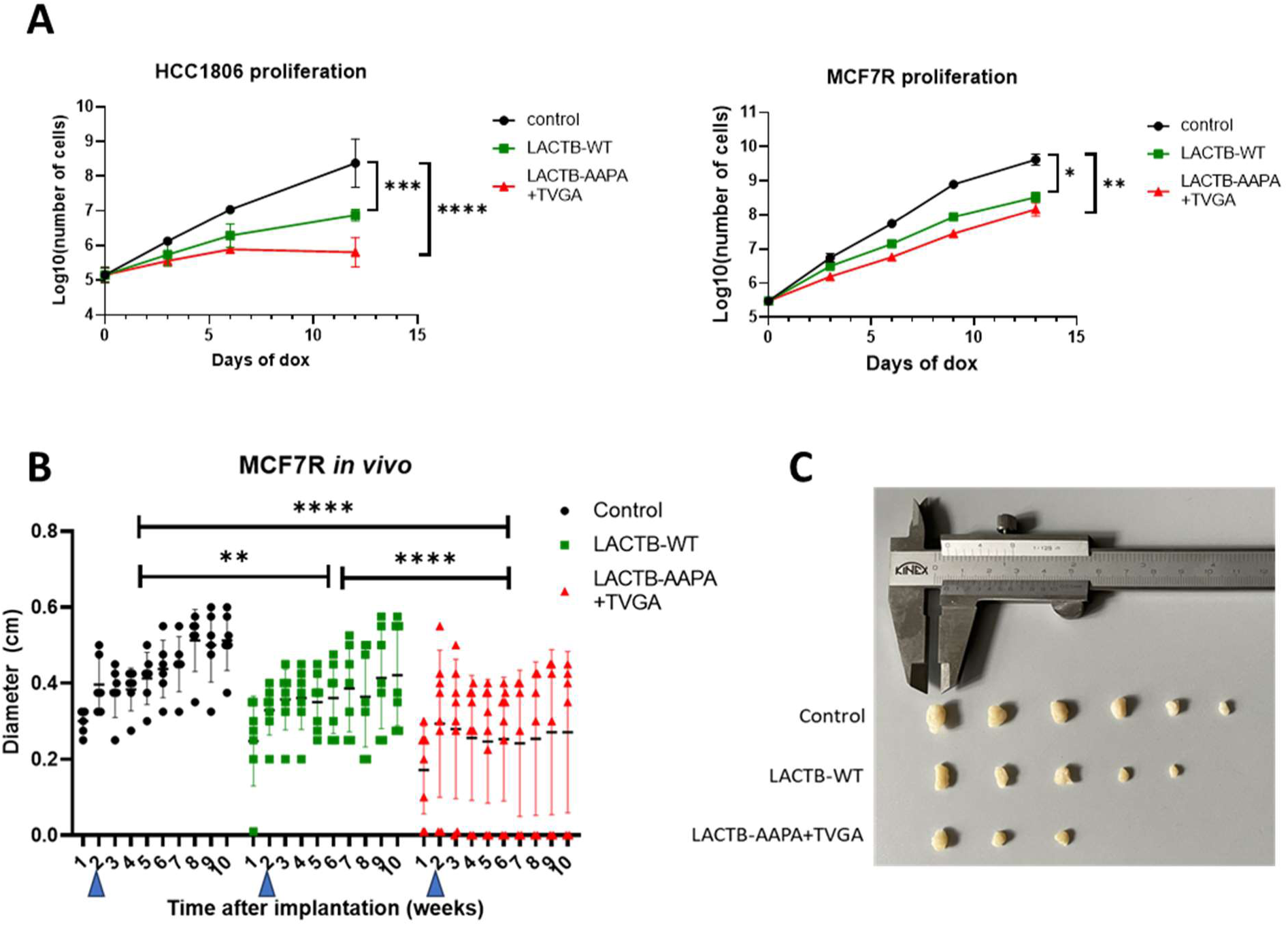
The autoproteolytic activity of LACTB influences its tumor suppressive ability *in vitro* and *in vivo*. **(A)** Proliferation curve of MCF7R and HCC1806 cells upon expression of dox-inducible LACTB-WT and LACTB-AAPA+TVGA. Cell numbers were determined by counting live cells every 3 days up to day 12. ns p>0.05; * p<0.05; ** p<0.01; *** p<0.001, **** p<0.0001. **(B)** Growth of tumors formed by injection of MCF7R (black), MCF7R LACTB-WT (green) and MCF7R LACTB-AAPA+TVGA (red) cells into the mouse mammary fat pad. Doxycycline treatment was initiated at 2 weeks of tumor growth (blue arrows). Statistics were calculated comparing the mean of each group using 2way-ANOVA test; ** p<0.01; **** p<0.0001. **(C)** Comparison of tumor sizes between MCF7R, MCF7R-LACTB-WT and MCF7R-LACTB-AAPA+TVGA. Some tumors in MCF7R-LACTB-AAPA+TVGA group were completely eliminated during the course of the doxycycline treatment.

### 4. MRPS34 is an interacting partner of LACTB

LACTB was shown by many studies to be efficiently downregulated in cancer cells [10,15,18,20,25], and our results presented above pointed to the possibility that LACTB might be downregulated by cancer cells through induction of its autoproteolytic activity. This prompted us to research this possible mode of regulation and to uncover the regulator/s of LACTB’s autoproteolytic activity in cancer cells using immunoprecipitation assays. In these experiments, we pulled down LACTB protein complexes from MCF7R cells expressing endogenous LACTB or doxycycline inducible wild type LACTB (1 day induction) and analyzed them by LC-MS/MS for the presence of LACTB’s binding partners. The proteins identified through mass spectrometry (Suppl. Data 2) were then validated for their ability to bind LACTB through secondary immunoprecipitation experiments and subsequent Western Blot analysis. The results showed that mitochondrial small ribosomal subunit protein mS34 (MRPS34) was efficiently pulled down through LACTB antibody (Fig. 4A). Within mitochondria, MRPS34 helps to stabilize the structure of the small ribosomal subunit and is essential for the efficient translation of mitochondrial-encoded genes. We did not observe any interaction of LACTB with other tested members of the MRPS-family, such as DAP3 (MRPS29) or MRPS22 (Suppl. Fig. 4A). The interaction of LACTB and MRPS34 proteins was further confirmed *in vitro* by subsequent dot blot experiments (Fig. 4B) and in cellular context by proximity ligation assay (PLA) in MCF7R cells (Fig. 4C-D). PLA was also performed using antibodies against other mitochondrial proteins (such as ATP5A and DAP3) which, in contrast to MRPS34, showed no interaction with LACTB (Fig. 4D). Using LZerD software we performed protein docking to predict the interaction between LACTB and MRPS34. The predicted models show binding of MRPS34 to LACTB tetramer and the formation of a salt bridge between Asp186(LACTB) and Arg101(MRPS34) (Suppl. Fig. 4B).

**Figure 4.**
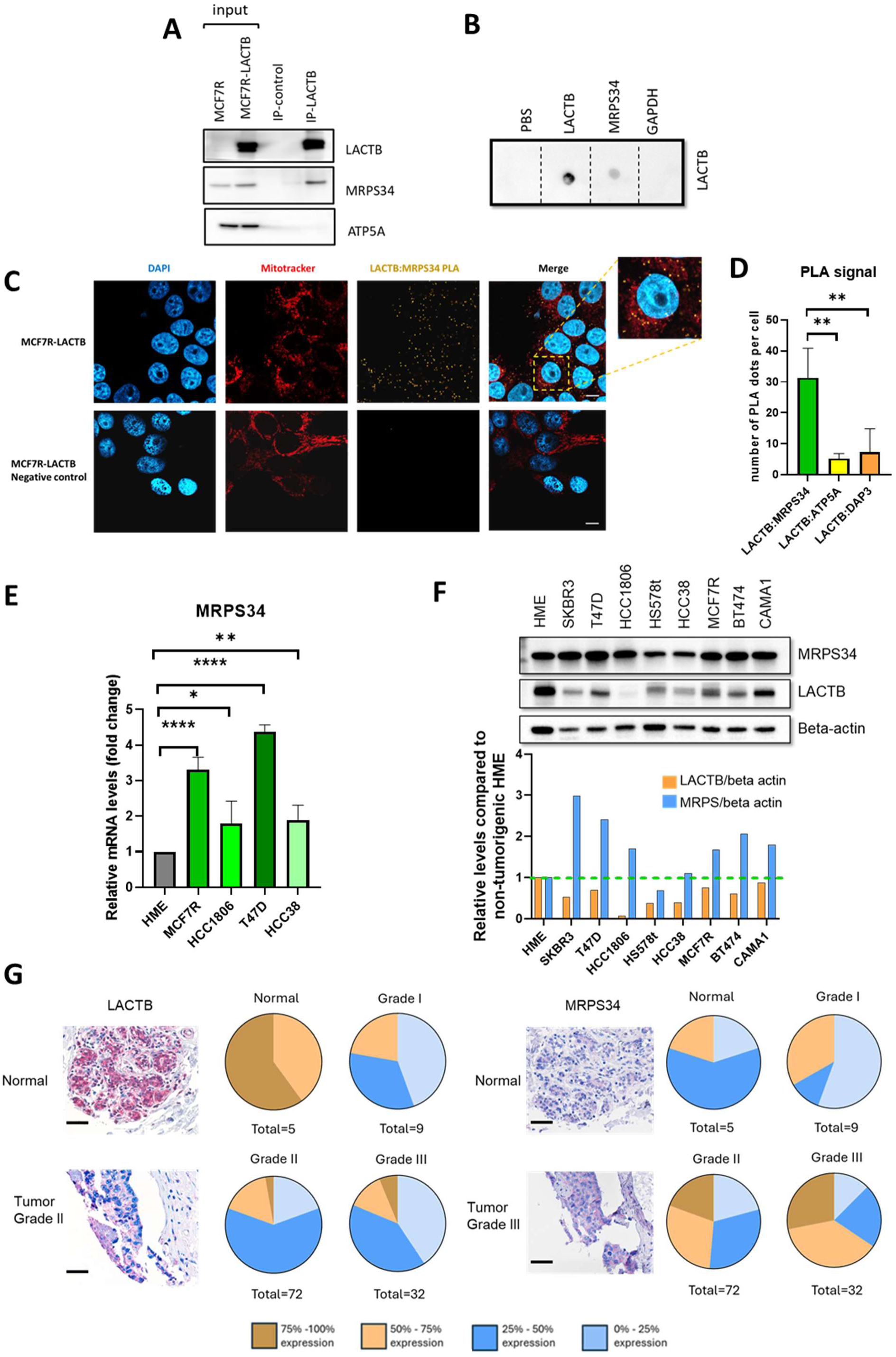
LACTB is in complex with MRPS34. **(A)** Immunoprecipitation (IP) experiments through LACTB antibody followed by Western blot analysis with antibodies against LACTB, MRPS34 and ATP5A. The input consists of lysate of MCF7R and MCF7R overexpressing doxycycline-inducible LACTB for 1 day. IP-control was done by immunoprecipitating with Resistin IgG antibody. **(B)** Dot blot assay with LACTB, MRPS34 and GAPDH recombinant proteins, followed by incubation with recombinant LACTB-WT and immunoblotting with LACTB antibody**. (C)** PLA assay performed in MCF7R-LACTB cells, using LACTB and MRPS34 primary antibodies. Pictures in the bottom (negative control) were processed without adding PLA probes. DAPI is depicted in blue, mitochondria are represented in red stained with MitoTracker Deep Red, and yellow is the positive PLA signal (dots represent *in situ* interaction). Scale bar is 10 µM. **(D)** Quantification of the number of PLA dots per cell. The experiment was performed in MCF7R cells, and it was used with the following combinations of antibodies: LACTB-MRPS34, LACTB-ATP5A and LACTB-DAP3. Statistics were performed using unpaired Student’s *t*-test. ** p<0.01. **(E)** mRNA expression of endogenous MRPS34, normalized to GAPDH, in non-tumorigenic HME, and in cancer cell lines MCF7R, HCC1806, HCC38 and T47D. Statistics were calculated using unpaired *t*-test, * p<0.05; ** p<0.01; **** p<0.0001. **(F)** Protein expression of endogenous MRPS34 and LACTB in a panel of breast cancer cells and non-tumorigenic HME cells. Graphs represent the quantification of the MRPS34 and LACTB intensity normalized to the beta-actin intensity for each cell line, normalized to non-tumorigenic HME cells. Representative experiment is shown. All experiments were performed three times. **(G)** Immunohistology staining of human breast clinical tissues (n=119) of normal or tumorigenic (grade 1, 2, 3) origin for LACTB and MRPS34 protein expression. Scale bar = 50 µm.

We next investigated the expression pattern of MRPS34 in normal and cancer cells. Bioinformatic analysis showed that tumor types (such as breast, lung, colon), where LACTB was previously reported to be downregulated [10,15,18,24], have a significantly higher expression of MRPS34 compared to normal tissues (Suppl. Fig. 4C); in breast cancer this upregulation occurs throughout all stages, subclasses and types (Suppl. Fig. 4D-F). Moreover, patients with higher expression of MRPS34 tend to have a lower survival rate (Suppl. Fig. 4G-H). RT-qPCR analysis confirmed that MRPS34 is upregulated at mRNA levels also in breast tumorigenic cell lines such MCF7R, HCC1806, T47D and HCC38 when compared to non-tumorigenic HME cells (Fig. 4E). We further profiled the protein expression pattern of endogenous MRPS34 and LACTB by western blot in a panel of 8 breast cancer cell lines and found that MRPS34 protein is upregulated and LACTB is downregulated in most of the tested cancer cell lines compared to their levels in HME cells (Fig. 4F). The RT-qPCR analysis of LACTB mRNA levels in several tumorigenic cell lines showed no strong correlation with its downregulation on a protein level suggesting that LACTB is, in these cell lines, downregulated on a post-transcriptional level (Suppl. Fig. 4I). Furthermore, we extended our experiments to human breast clinical tissues (n=119) of normal or tumorigenic (grade 1, 2, 3) origin. Immunohistology staining of primary tissues showed that tissues where LACTB protein was downregulated (tumorigenic tissues) had increased expression of MRPS34 (Fig. 4G). Thus, all these sets of experiments showed interaction between LACTB and MRPS34 and pointed to the inverse expression pattern of LACTB and MRPS34 in cell lines and primary human tissues suggesting the clinical significance of these findings.

### 5. MRPS34 is a negative regulator of LACTB through enhancement of its autoproteolytic activity

To further investigate the role of MRPS34 in LACTB regulation we employed shRNA against MRPS34 in a panel of tumorigenic breast cancer cell lines (MCF7R, T47D, HS578t, HCC1806 and HCC38) and monitored the levels of endogenous LACTB upon MRPS34 downregulation. We observed that downregulation of MRPS34 led to increases in endogenous LACTB protein levels in majority of the tumorigenic cell lines studied, revealing MRPS34’s role in negatively regulating LACTB levels in breast cancer cells (Fig. 5A). Of note, strong MRPS34 knockdown had a toxic effect on several cancer cell lines, therefore we aimed for partial knockdown of MRPS34. Even under these conditions, significant increases in LACTB expression levels were readily observed. We next overexpressed MRPS34 protein in MCF7R, T47D, HCC1806, HS578t and HCC38, and noticed robust decreases of endogenous LACTB levels in all cell lines tested (Fig. 5B). We did not observe any difference in LACTB mRNA levels in MCF7R, HCC38 and T47D cells overexpressing MRPS34 when compared to the control cells, suggesting that the negative regulation of LACTB protein levels by MRPS34 occurs at a post-transcriptional level (Suppl. Fig. 5A). Since our previous results showed that LACTB can degrade itself through autoproteolytic activity and that MRPS34 is a negative regulator of LACTB, we investigated the possibility whether the MRPS34 exerts its negative regulation of LACTB by enhancing its autoproteolytic activity. We therefore incubated recombinant LACTB-WT with and without MRPS34 for different time points, from 18 hours to 48 hours. MRPS34’s presence significantly accelerated the autoproteolytic activity of LACTB, which was observable at early time points of co-incubation (Fig. 5C). Of note, we noticed that the autoproteolytic activity of LACTB in the presence of MRPS34 was more apparent and appeared at earlier timepoints in the multimeric LACTB forms. In the presence of the inhibitor of the enzymatic activity of LACTB, no autoproteolytic activity was observed at any of the time points (Fig. 5C). Regarding the enzymatic activity efficiency directed against peptide probe, recombinant LACTB incubated together with MRPS34 displayed a decrease in the enzymatic activity (Suppl. Fig. 5B), confirming the onset of the accelerated degradation of LACTB-WT in the presence of MRPS34. However, when recombinant LACTB was incubated together with other mitochondrial proteins (COQ6, HSBP1, TOM20, HIGD1A, ASCL3) there were no decreases in its enzymatic activity (Suppl. Fig. 5B) confirming the specific effect MRPS34 has on LACTB biology. We next assessed whether MRPS34 overexpression could affect the proliferation rate of breast cancer cells with concomitant LACTB expression. The results showed that LACTB-induced decrease of cell proliferation in MCF7R and HS578t cells was indeed partially recovered by concomitant MRPS34 induction (Fig. 5D and Suppl. Fig. 5C), confirming the negative regulation of LACTB by MRPS34 and the resulting positive effect on the viability of cancer cells upon MRPS34 induction. Similarly, flow cytometry analysis examining the cell cycle progression in MCF7R cells showed that overexpression of MRPS34 in LACTB-induced cells partly restores G1 cell cycle progression and significantly increased S phase content when compared to LACTB-induced cells without MRPS34 (Suppl. Fig. 5D) further confirming that expression of MRPS34 counteracts the tumor suppressive effects of LACTB.

**Figure 5.**
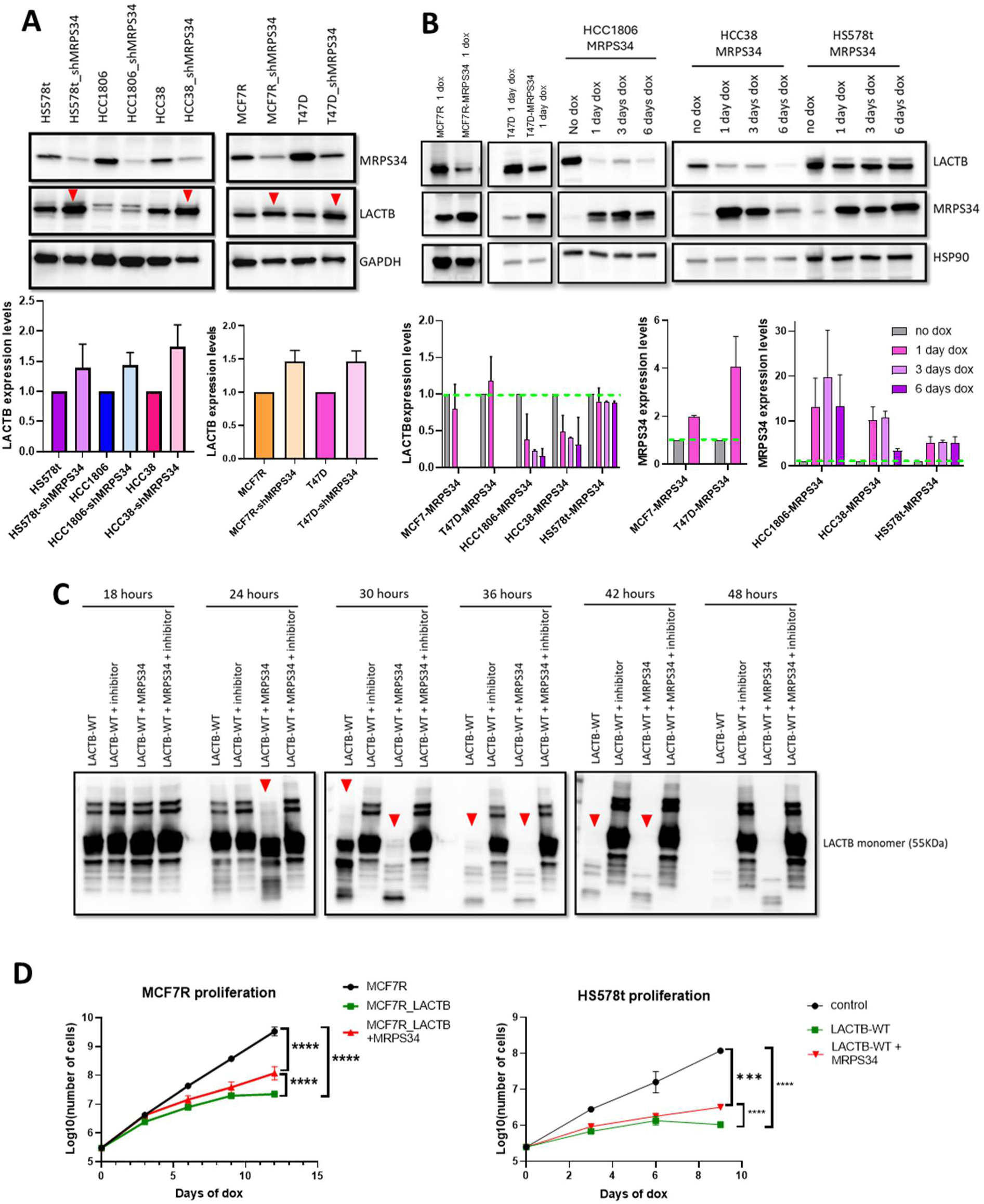
MRPS34 is a negative regulator of LACTB. **(A)** LACTB protein expression levels in control cells and upon downregulation of MRPS34 in a panel of tumorigenic (MCF7R, T47D, HS578t, HCC1806, HCC38) cell lines. Graphs represent the quantification of the monomer LACTB intensity normalized to the GAPDH intensity for each cell line, followed by normalization of shMRPS34 to the respective control cells. The experiment was performed three times. **(B)** LACTB expression levels in control cells and upon overexpression of dox-inducible MRPS34, in a panel of tumorigenic (MCF7R, T47D, HCC1806, HS578t, HCC38) cell lines. Graphs represent the quantification of MRPS34 or of the monomer LACTB intensity normalized to the GAPDH (in MCF7R) or HSP90 (in T47D, HCC1806, HCC38 and HS578t) intensity for each cell line. The green perforated line shows the level of MRPS34 or LACTB in control cells. **(C)** LACTB and LACTB + MRPS34 recombinant proteins, with and without LACTB inhibitor, were incubated at 25 °C for different time points from 18 hours to 48 hours in H_2_O, followed by Western blot and immunoblotting with LACTB antibody. **(D)** Proliferation rate of MCF7R and HS578t cells with and without dox-inducible LACTB and MRPS34, upon doxycycline induction for up to 12 days. The number of cells is shown on a logarithmic scale. The experiment was performed 3 times for MCF7R cells and 2 times for HS578t cells. *** p<0.001; **** p<0.0001

Overall, these results showed that MRPS34 is a negative regulator of LACTB capable of increasing LACTB’s autoproteolytic activity thus enhancing cancer cell survival.

## Discussion

In this study, we uncovered the mechanisms cancer cells use to post-translationally downregulate LACTB tumor suppressor thus keeping the expression levels of LACTB low. Through our search for a LACTB substrate we showed that LACTB possesses autoproteolytic activity that is enhanced by mitochondrial MRPS34 protein, which, through interaction with LACTB, acts as a negative regulator of LACTB. The LACTB-MRPS34 axis is thrown out of balance by cancer cells, which, through upregulation of MRPS34, achieve efficient down-regulation of LACTB, realized through its own auto-destruction (Fig. 6).

**Figure 6.**
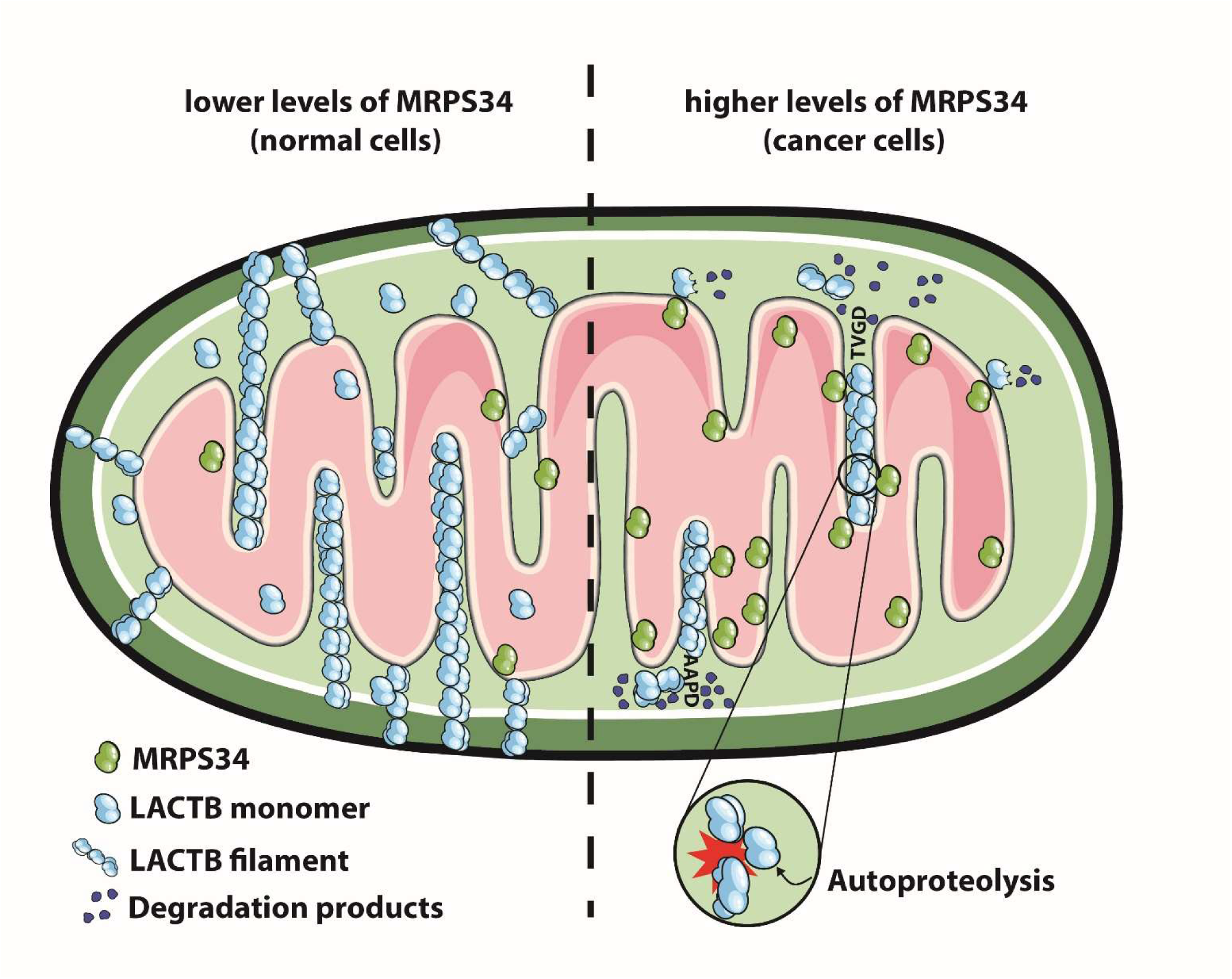
Graphic representation of LACTB’s mechanism of regulation in breast cancer cells. In normal non-tumorigenic cells, the levels of MRPS34 are low and LACTB is present in abundance in the form of monomers and/or multimers/filaments mainly in the intermembrane mitochondrial space. In cancer cells the levels of MRPS34 are upregulated, MRPS34 interacts with LACTB what leads to onset of its autoproteolytic activity. This activity is primarily realized against AAPD and TVGD sequences within LACTB and leads to self-degradation of LACTB resulting in lower amounts of LACTB in cancer cells.

Mammalian mitochondria contain mitoribosomes which consist of a small (28S) and large (39S) subunits, both containing mtDNA-encoded rRNA – mainly located in the interior of the mitoribosome, and nuclear-encoded mitochondrial ribosomal proteins (MRPs) – present on the mitoribosome surface [33–35]. The development of many cancers has been associated with enhanced expression of MRPs, what could be explained by the increased metabolic activity of cancer cells [36]. LACTB has been previously associated with the large subunit of the mammalian mitochondrial ribosome [37] and our study shows its ability to also interact with MRPS34, the small subunit of the mitoribosome, which is localized in the matrix of mitochondria tightly tethered to inner mitochondrial membranes [38]. Later studies showed that LACTB is also localized in the mitochondrial intermembrane space, where it forms filaments that are suggested to be embedded in the inner mitochondrial membrane [39]. Therefore, it is plausible for LACTB and MRPS34 to interact in the mitochondrial matrix or within the inner mitochondrial membrane area.

We speculate that under normal physiological conditions, the autoproteolytic activity of LACTB serves as a physiological regulatory mechanism of mitochondrial LACTB protein levels or its maturation stages. LACTB, through its ability to form filaments within the intermembrane mitochondrial space, was suggested to play a role in the structural organization of mitochondria. It is possible that the autoproteolytic processing of LACTB might serve, through increased/decreased ratios of LACTB monomers and LACTB filaments, as a fast and efficient mitochondrial re-structuring and adaptation mechanism to various cellular stimuli (modulation of differentiation, mitochondrial pore permeability, metabolism, ions accessibility or signaling pathways stemming from mitochondria). Our previous work [40] showed sensitivity of enzymatic activities of LACTB to different pH, buffers, salts and ions suggesting that LACTB might serve either as a sensor of these conditions within mitochondria or might be regulated by them to fulfill distinct mitochondrial functions. Depending on the nature of these conditions LACTB, through switching between the monomeric and/or filamentous form or between enzymatic activity directed against cellular substrate or itself, influences the structure of the mitochondria and the processes realized within the mitochondria.

Overall, our study discovered novel aspects of tumor suppressor regulation by cancer cells and uncovered additional vulnerabilities of cancer cells which might help us in our fight against cancer progression.

## Supporting information

PICS assay

Binding partners

## Acknowledgements

This work was supported by the Czech Science Foundation (18-24473Y), EMBO Installation Grant and Ministry of Education, Youth and Sports, and the European Social Fund; OP RDE; Project: “IOCB MSCA Mobility III” (No. CZ.02.2.69/0.0/0.0/19_074/0016322) (Z.K). Additional grants were provided by the project National Institute for Cancer Research (Programme EXCELES, ID Project No. LX22NPO5102) - Funded by the European Union - Next Generation EU (Z.K), the Ministry of Science and Technology of the People’s Republic of China (2020YFA0803300, Y.-X.F.), the National Natural Science Foundation of China (32270783, Y.-X.F.). Mouse experiments were performed by Petra Kralova Viziova and supported by the Czech Academy of Sciences RVO 68378050, LM2018126 and LM2023036 Czech Centre for Phenogenomics provided by MEYS and by CZ.02.1.01/0.0/0.0/18_046/0015861 CCP Infrastructure Upgrade II by MEYS and ESIF. Proteomics LC/MS analyses were performed by Karel Harant in Laboratory of Mass Spectrometry at Biocev research center, Faculty of Science, Charles University.

## Contributions

S.E.L. designed and performed the experiments, analyzed the data and prepared the manuscript. J.G.M performed the FACS, and clinical tissue experiments and analyzed the data. J.F.M., P.M., G.R.G, CH.W.P., P.J., S.M., Y.F. helped with experiment design. A.B., M.S performed the modified PICS assay, S.K.M. performed the bioinformatics interaction studies, and B.M. provided technical support. Z.K. designed and supervised this study and prepared the manuscript.

---

**Supplementary Figure 1.**
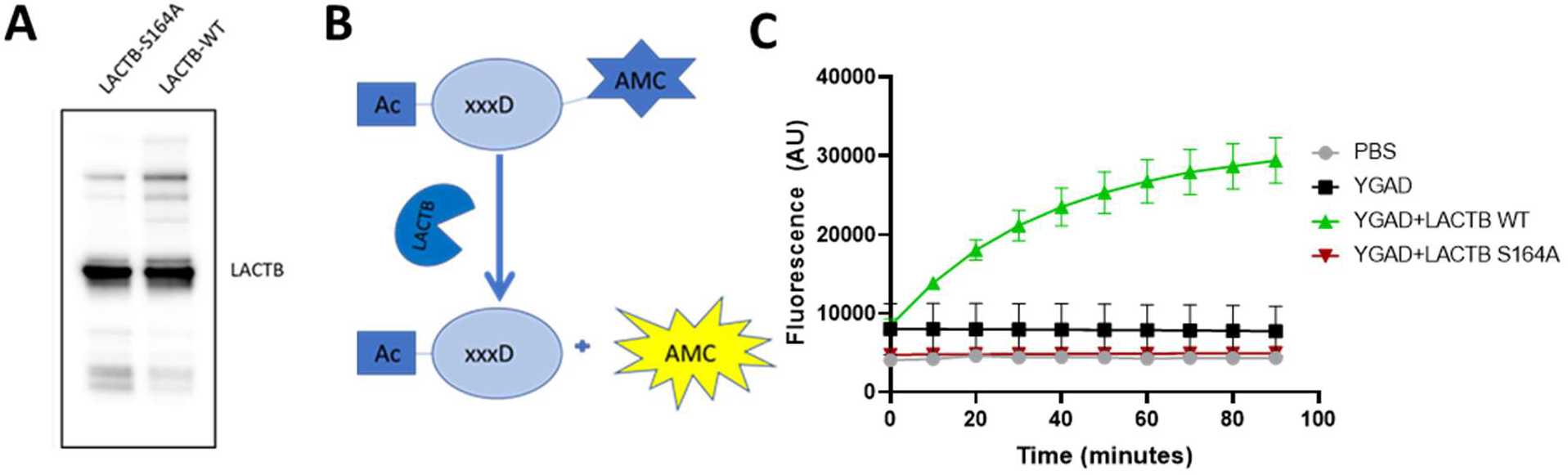
**(A)** The expression level of LACTB-WT and LACTB-S164A recombinant proteins shown by Western blot analysis. **(B)** Graphical illustration of LACTB’s enzymatic activity assay using tetrapeptides (such as YVAD and YGAD) linked to AMC fluorescent probe. Upon cleavage of the probe by LACTB the AMC moiety is released and emits fluorescence. **(C)** Comparison of the cleavage ability of LACTB-WT and LACTB-S164A catalytic mutant against the YGAD probe.

**Supplementary Figure 2.**
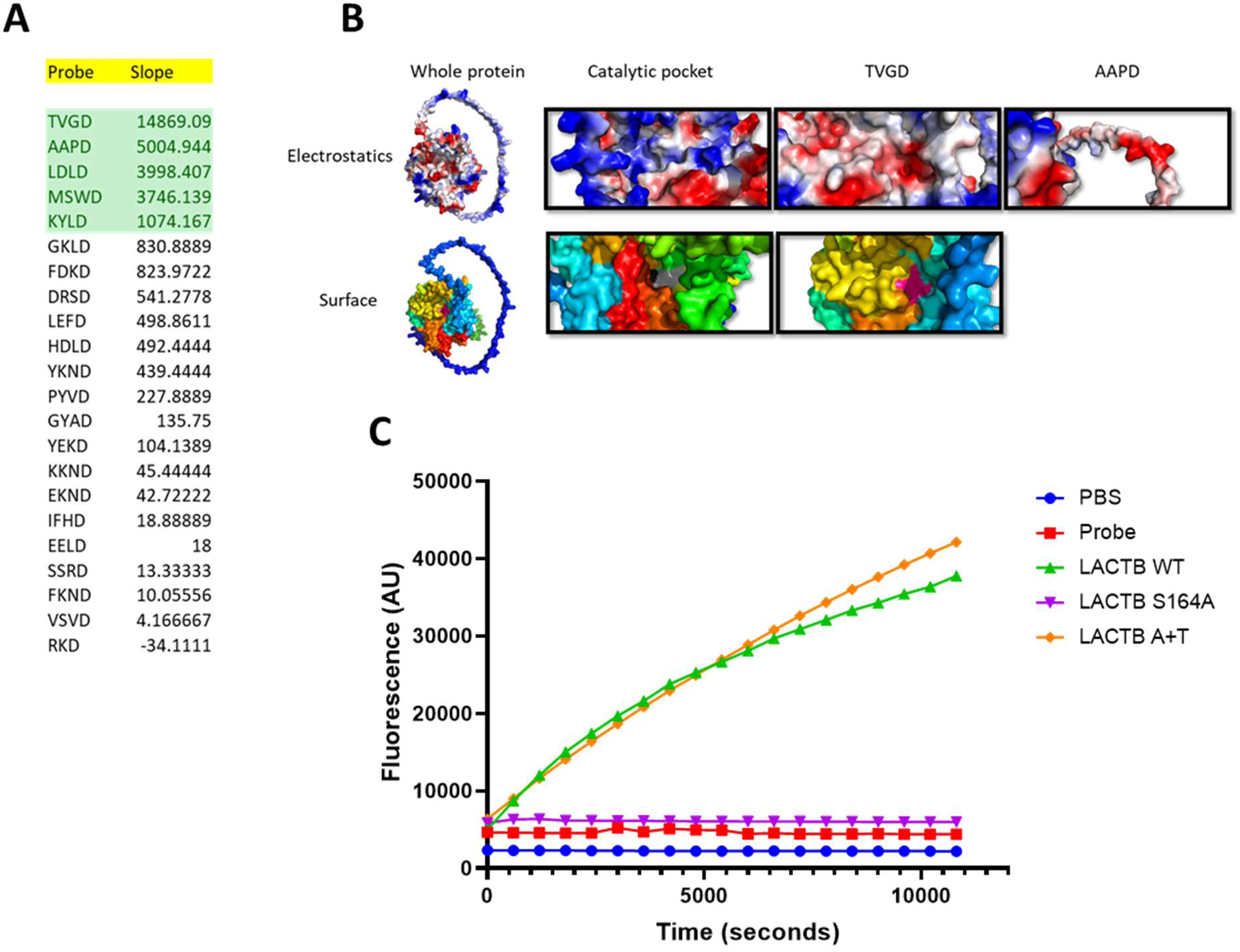
**(A)** The list of peptides, fused to AMC fluorescent probe, representing all tetrapeptides ending in aspartic acid within the LACTB sequence that were tested for their ability to be cleaved by LACTB. Highlighted in the table are the peptides most efficiently cleaved by LACTB, as calculated by subtracting the first fluorescence measurement to the last fluorescence measurement (slope). **(B)** Upper panel: electrostatic comparison between the catalytic pocket and TVGD and AAPD sites; Bottom panel: surface analysis of full length LACTB-WT, showing the catalytic pocket (grey) and TVGD (magenta) regions. **(C)** Enzymatic activity of the autoproteolytic mutant LACTB-AAPA+TVGA (A+T) compared to the catalytically inactive mutant S164A and LACTB WT. Experiment was performed twice.

**Supplementary Figure 3.**
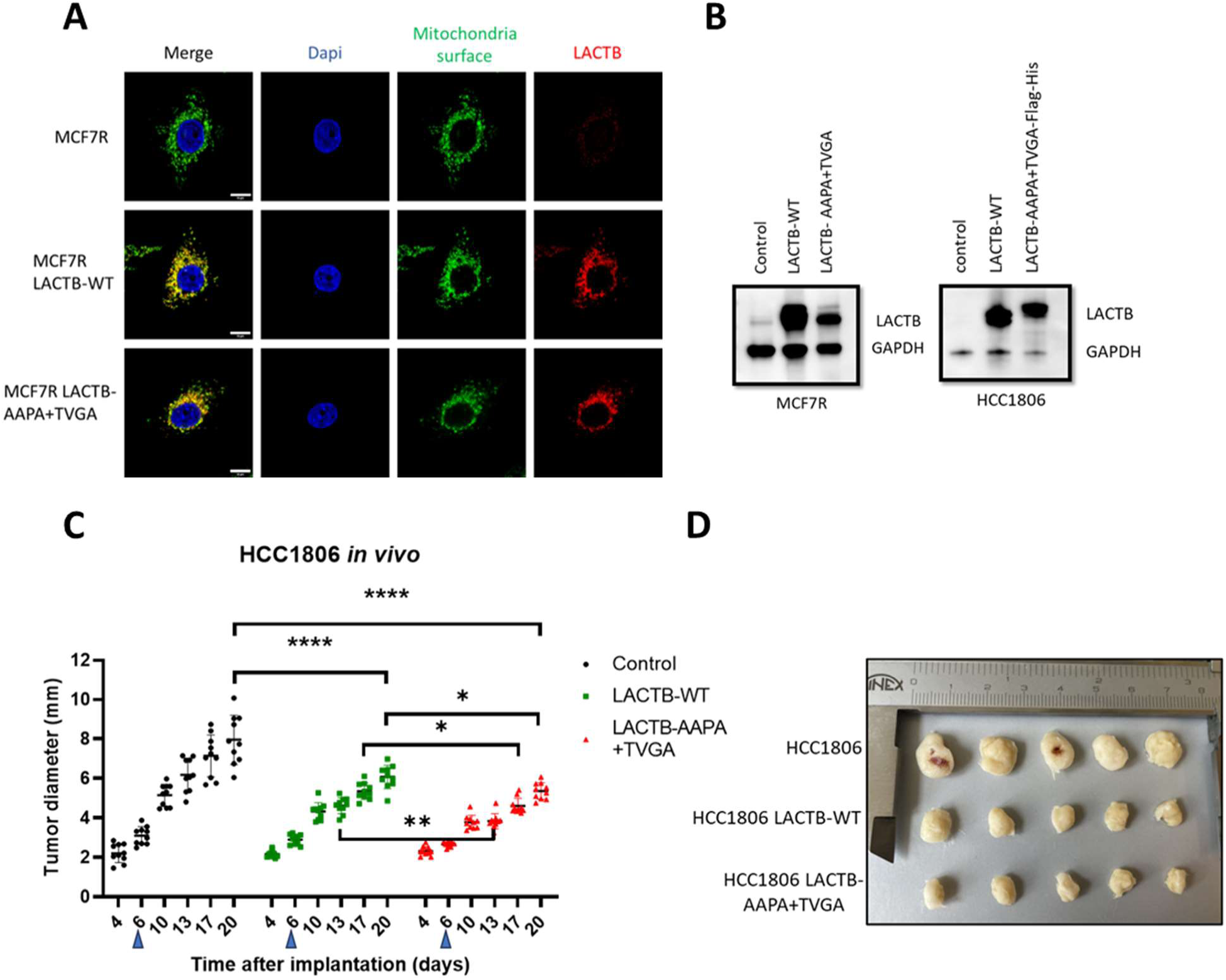
**(A)** Confocal microscopy images of the localization of LACTB-WT and LACTB-AAPA+TVGA in MCF7R, after one day of induction. Blue corresponds to DAPI, green labels mitochondria surface (Merck, MAB1273) and red represents LACTB. Scale bar is 5 µm. **(B)** Expression levels of LACTB-WT and LACTB-AAPA+TVGA in MCF7R and HCC1806 cells. **(C)** Growth of tumors formed by injection of control HCC1806 (black), HCC1806 LACTB-WT (green) and HCC1806 LACTB-AAPA+TVGA (red) cells into the mouse mammary fat pad. Doxycycline treatment was initiated at 6 days of tumor growth (blue arrows). Statistics were calculated using 2way-ANOVA, * p<0.05; ** p<0.01, **** p<0.0001. **(D)** Comparison of tumor sizes between HCC1806, HCC1806-LACTB-WT and HCC1806-LACTB-AAPA+TVGA

**Supplementary Figure 4.**
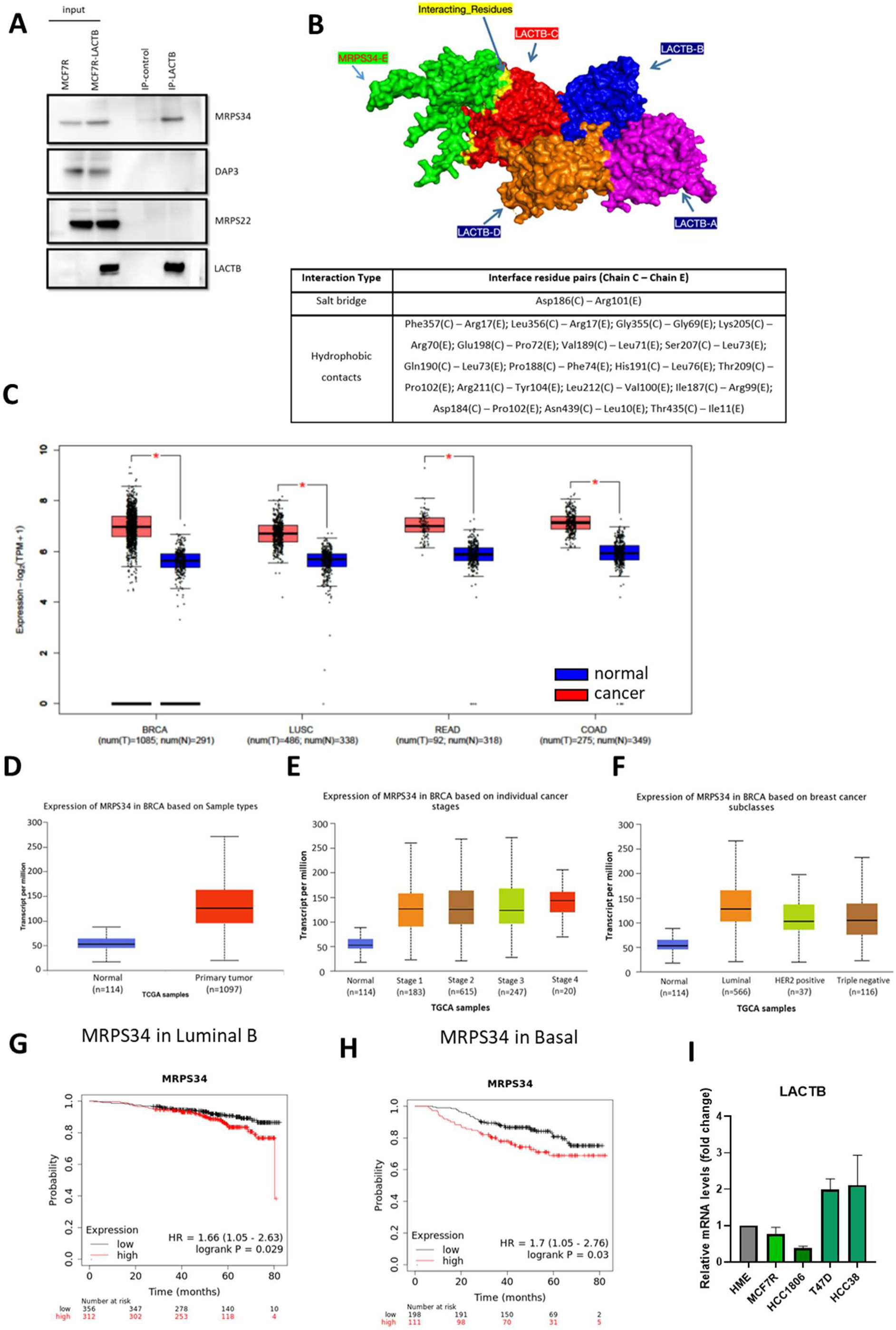
**(A)** Immunoprecipitation (IP) through LACTB antibody followed by Western blot with antibodies against MRPS34, DAP3/MRPS29 and MRPS22. The input consists of lysate of MCF7R and MCF7R overexpressing doxycycline-inducible LACTB for 1 day. IP-control was done by immunoprecipitating with Resistin IgG antibody. Part of this figure is shared with Figure 4A. **(B)** Structural depiction of the interaction between the LACTB tetramer and MRPS34. The LACTB subunits (chains A–D) are illustrated in magenta, blue, orange, and red, respectively, while MRPS34 (chain E) is shown in green. Residues involved in interfacial contacts are highlighted in yellow. The accompanying table details the interface interactions, categorizing residue pairs into salt bridges, and hydrophobic contacts. **(C)** Expression of MRPS34 in normal (blue) and primary cancer (red) tissues of breast, lung, rectum and colon cancers. Graph was obtained using GEPIA2 tool. **(D-F)** Expression of MRPS34 in normal and breast cancer (BRCA) tissues stratified by different stages and subtypes. Graph obtained using UALCAN tool. **(G)** Effects of the expression of MRPS34 on overall survival in patients with luminal B breast cancer, using KMplotter tool. **(H)** Effects of the expression of MRPS34 on overall survival in patients with basal breast cancer, using KMplotter tool.

**Supplementary Figure 5.**
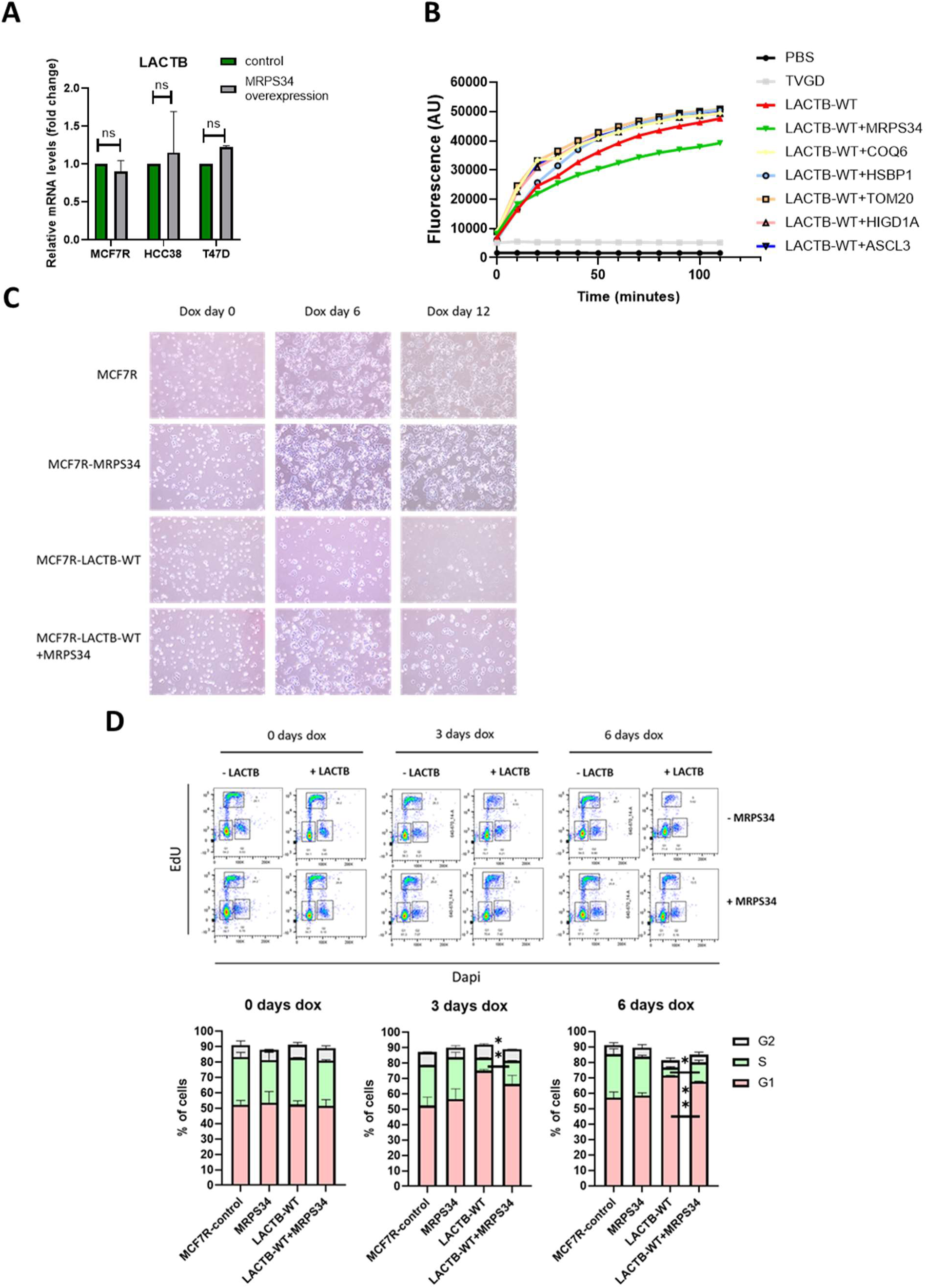
**(A)** Endogenous LACTB mRNA levels in MCF7R, HCC38 and T47D cancer cells with or without MRPS34 induction for 1 day, normalized to GAPDH. Experiment performed twice. **(B)** Enzymatic activity of LACTB directed against a peptide probe in the presence of MRPS34 and other mitochondrial proteins. LACTB-WT was pre-incubated with the respective proteins for 48 hours at 25 °C prior to the assessment of its enzymatic activity. **(C)** Light microscopy images of MCF7R cells, expressing LACTB, MRPS34 or both for 0, 6 and 12 days. **(D)** Flow cytometry-based cell cycle analysis of EdU/DAPI in MCF7R cells. Comparison of the DNA synthesis (S-phase) and G1 phase in MCF7R-LACTB-WT and MCF7R-LACTB-WT+MRPS34 cells upon doxycycline induction (day 0, 3 and 6), using Student’s *t*-test. Experiments were performed three times. Significance was reached when P < 0.05. * p<0.05; ** p<0.01.

**Supplementary table 1.**

| **Cell line** | **Media** | **Source** | **RRIDs** |
| --- | --- | --- | --- |
| HEK293T | DMEM + 10 % FBS + 1 % P/S | ATCC | CVCL_0063 |
| MCF7/R | DMEM + 10 % FBS + 1 % P/S | Merck | CVCL_0031 |
| HMEC/HME | MEGM (Lonza) | Lonza | CVCL_3383 |
| HCC1806 | RPMI + 10 % FBS + 1 % P/S | ATCC | CVCL_1258 |
| HCC38 | RPMI + 10 % FBS + 1 % P/S | ATCC | CVCL_1267 |
| HS578t | RPMI + 10 % FBS + 1 % P/S | ATCC | CVCL_0332 |
| T47D | RPMI + 10 % FBS + 1 % P/S | ATCC | CVCL_0553 |
| SKBR3 | DMEM + 10 % IFS + 1 % P/S | ATCC | CVCL_0033 |
| CAMA1 | DMEM + 10 % IFS + 1 % P/S | ATCC | CVCL_1115 |
| BT474 | RPMI + 10 % FBS + 1 % P/S | ATCC | CVCL_0179 |

**Supplementary table 2.**
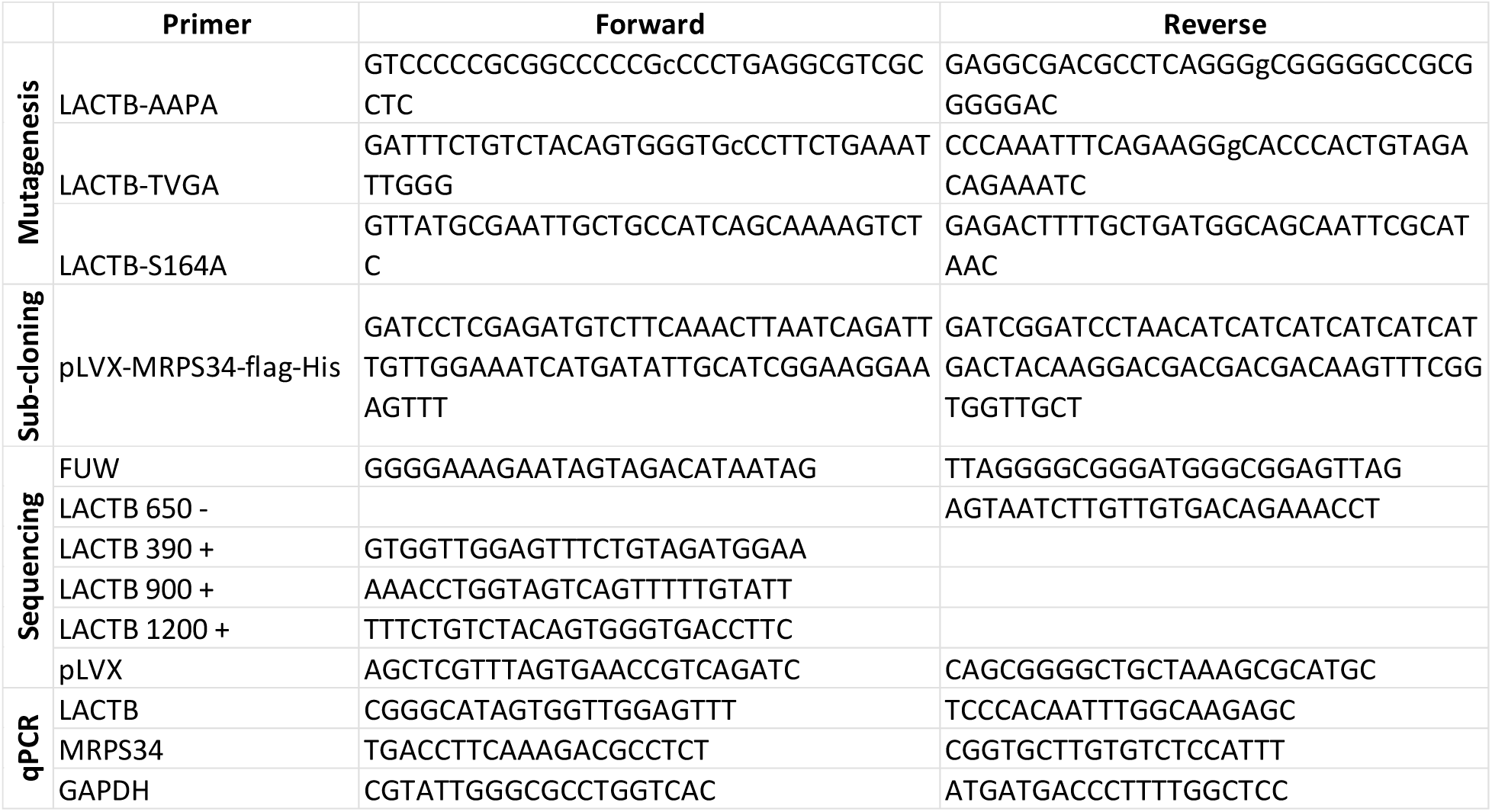

**Supplementary table 3.**
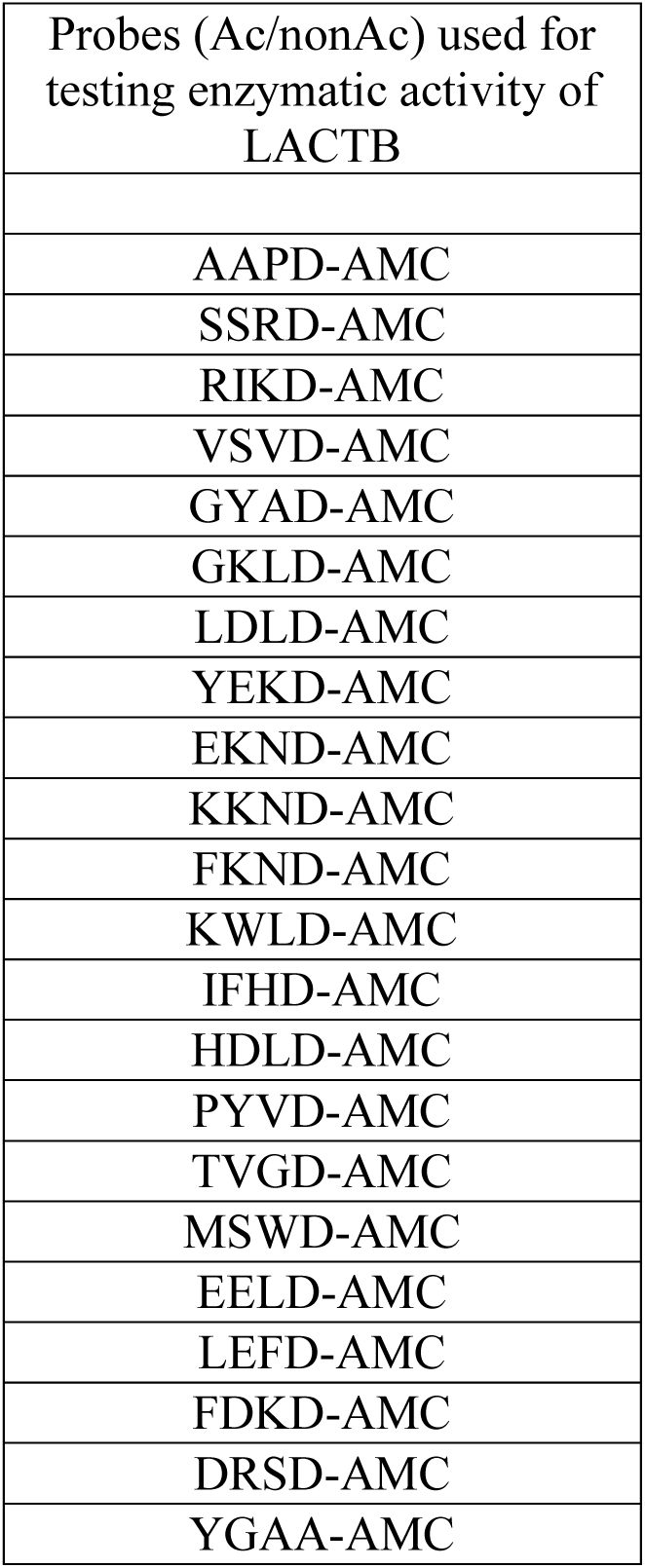

**Supplementary table 4.**

| Antibodies and reagents | Dilution or concentration | Catalog # |
| --- | --- | --- |
| ATP5A | WB: 1000X | AB14748 |
| beta-actin | WB: 1000X | 66009-1-Ig |
| DAP3 | WB: 1000X | 10276-1-AP |
| GAPDH | WB: 1000X | 60004-1-Ig |
| HSP90 | WB: 3000X | 13171-1-AP |
| LACTB-rabbit | IHC: 250X | PAJ512Hu01 |
| LACTB-rabbit | WB: 1000X, IF: 250X | 18195-1-AP |
| Mitochondria, surface of intact mitochondria clone 113-1 | IF: 250X | MAB1273 |
| MRPS22 | WB: 1000X | 10984-1-AP |
| MRPS34-rabbit | WB: 500X, IF: 200X, IHC: 250X | 15166-1-AP |
| MitoTracker Deep Red | IF: 0.25 μM | M22426 |

